# Common and unique neurophysiological signatures for the stopping and revising of actions reveal the temporal dynamics of inhibitory control

**DOI:** 10.1101/2024.06.18.597172

**Authors:** Mario Hervault, Jan R. Wessel

## Abstract

Inhibitory control is a crucial cognitive-control ability for behavioral flexibility that has been extensively investigated through action-stopping tasks. Multiple neurophysiological features have been proposed to represent ‘signatures’ of inhibitory control during action-stopping, though the processes signified by these signatures are still controversially discussed. The present study aimed to disentangle these processes by comparing simple stopping situations with those in which additional action revisions were needed. Three experiments in female and male humans were performed to characterize the neurophysiological dynamics involved in action-stopping and - changing, with hypotheses derived from recently developed two-stage ‘pause-then-cancel’ models of inhibitory control. Both stopping and revising an action triggered an early broad ‘pause’-process, marked by frontal EEG β-bursts and non-selective suppression of corticospinal excitability. However, partial-EMG responses showed that motor activity was only partially inhibited by this ‘pause’, and that this activity can be further modulated during action-revision. In line with two-stage models of inhibitory control, subsequent frontocentral EEG activity after this initial ‘pause’ selectively scaled depending on the required action revisions, with more activity observed for more complex revisions. This demonstrates the presence of a selective, effector-specific ‘retune’ phase as the second process involved in action-stopping and -revision. Together, these findings show that inhibitory control is implemented over an extended period of time and in at least two phases. We are further able to align the most commonly proposed neurophysiological signatures to these phases and show that they are differentially modulated by the complexity of action-revision.

**Significance Statement:** Inhibitory control is one of the most important control processes by which humans can regulate their behavior. Multiple neurophysiological signatures have been proposed to reflect inhibitory control. However, these play out on different time scales and appear to reflect different aspects of cognitive control, which are controversially debated.

Recent two-stage models of inhibitory control have proposed that two phases implement the revisions of actions: ‘pause’ and ‘retune’. Here, we provide the first empirical evidence for this proposition: Action revisions engendered a common initial low-latency ‘pause’, during which motor activity is broadly suppressed. Later activity, however, distinguishes between simple stopping of actions and more complex action revisions. These findings provide novel insights into the sequential dynamics of human action control.

## Introduction

Inhibitory control is a core cognitive control ability that permits the revision of an already-initiated action, for example during rapid environmental changes. Over the past decades, its neural basis and clinical relevance have been extensively investigated based on action-stopping tasks (Logan and Cowan, 1984; Verbruggen et al., 2019). Multiple neurophysiological signatures of inhibitory control during action-stopping have been proposed (e.g., Huster et al., 2013; Wessel and Aron, 2017; Jana et al., 2020). They include the reduction of residual EMG activity on successfully stopped actions (partial-response EMG; prEMG) (Raud et al., 2022), corticospinal excitability (CSE) suppression (Duque et al., 2017), frontocentral β-bursts (Wessel, 2020), the P3 ERP (Wessel and Aron, 2015), and frontocentral θ power (Messel et al., 2021). It has been a major challenge to reconcile these signatures under a common theoretical framework, especially since they occur at different times after the signal to stop. Moreover, several signatures have been challenged with regard to their relationship to inhibitory control altogether (e.g., Huster et al., 2020; Skippen et al., 2020).

To account for this inconsistency, recently proposed ‘pause-then-cancel’ (PTC) models (Schmidt and Berke, 2017; Diesburg and Wessel, 2021) have conceptualized inhibitory control as a two-stage dynamic. An early ‘pause’ process is triggered by any unexpected signal to implement quick (∼150 ms) and broad inhibition of the motor system. This process has been proposed to be reflected in non-selective CSE suppression and the early suppression of prEMG (Tatz et al., 2021; Wessel et al., 2022). Additionally, β-bursts dynamics have been proposed to reflect subcortical-cortical communication (Diesburg et al., 2021) in a time frame corresponding to the ‘pause’ stage (Hannah et al., 2020; Tatz et al., 2023; Wadsley et al., 2023a). This ‘pause’ ostensibly buys time for a second process (‘cancel’) to stop the action in a selective fashion that depends on the task requirement (Waller et al., 2019; Tatz et al., 2021). Frontocentral P3 ERP and θ power increase have been proposed to index this process (Diesburg and Wessel, 2021). In addition, it has been proposed that the ‘cancel’ stage reflects a stop-specific instantiation of a more general ‘retuning’ process. In other words, ‘cancel’ only means ‘cancel’ if the task requires a complete cessation of all responding. If more complex adjustments are necessary, these ‘cancel’-process signatures should parametrically scale with the complexity of the response revision, and should reflect a ‘retuning’ of the response, rather than its cancellation (Diesburg and Wessel, 2021).

The current study aimed to comprehensively investigate these predictions regarding the purported neurophysiological signatures of inhibitory control under the human PTC model. We designed a series of tasks in which actions had to be revised with different degrees of complexity. Based on the PTC-model, we predicted that markers of the ‘pause’ stage (CSE suppression, prEMG, and frontal β-bursts) would be shared between all action-stopping and -changing requirements. Crucially, however, we predicted that purported markers of the ‘cancel’ stage (frontocentral P3 and θ power) would scale with the required response-revision (stopping, changing one response dimension, changing two response dimensions). This would confirm a key theoretical proposal of a more generic ‘pause-then-retune’ dynamic for action-revision (Diesburg and Wessel, 2021).

Alternatives to these hypotheses are at least twofold. First, action-changing might engage inhibitory processing more selectively than action-stopping (De Jong et al., 1995; Antons et al., 2019). If so, ‘pause’-stage signatures should not be observed (or be reduced) during action-changing. Second, previous fMRI studies (Kenner et al., 2010; Boecker et al., 2011) have identified that cortical blood-oxygen-level-dependent (BOLD) responses in action-changing was similar to the sum of the BOLD responses in action-stopping and action-initiation. This suggested that action-changing was performed by the sum of a generic stopping process plus a new response triggering (Boecker et al., 2013). If so, the ‘cancel’ stage purported signatures should index a similar inhibition dynamic across stopping and changing.

Together, this comprehensive investigation aims to test the validity of the recent neurocognitive models of inhibitory control, address contradictory unitary/dissociated views of stopping and changing processing (Krämer et al., 2011; Boecker et al., 2013; Rangel-Gomez et al., 2015; Elchlepp and Verbruggen, 2017), and understand specific clinical impairments in action-changing (McClure et al., 2005; Roberts and Husain, 2015; van den Wildenberg et al., 2017).

## Experiment 1

### Method

#### Hypothesis

Our main hypothesis for Experiment 1 was that both action-stopping and action-changing would induce short-latency corticospinal excitability (CSE) suppression, suggesting that a non-selective ‘pause’ inhibition dynamic is engaged in both situations. Stopping and changing have been investigated in mixed task-blocks (Boecker et al., 2007, 2011; Drueke et al., 2010; Rangel-Gomez et al., 2015). However, by intermixing STOP and STOP-CHANGE trials, one could bias the STOP-CHANGE condition towards exhibiting the same non-selective inhibition found during stopping, as participants may stop all responses first while they discern between stop- and change-signals (Bissett and Logan, 2014; Giarrocco et al., 2021). To allow a stronger test of the hypothesis that action-stopping and -changing would share this initial non-selective inhibitory phase, we hence tested them in separate blocks, allowing participants to use their preferred strategy for either behavior.

#### Participants

24 healthy individuals (15 females, mean age 19 years, SD = 1.6) participated in the experiment in exchange for course credit or an hourly payment of $15. They completed TMS screening questions (Rossi et al., 2011) and provided written informed consent. Procedures were approved by the local Institutional Review Board (IRB #201612707) and conducted in accordance with the Declaration of Helsinki.

In the absence of previous studies investigating corticospinal excitability in the STOP-CHANGE-signal task, the a priori calculation of our sample size was based on the behavioral comparison of STOP-signal and STOP-CHANGE-signal tasks. Boecker et al. reviewed 10 experiments comparing STOP-signal vs. STOP-CHANGE -signal reaction times (SSRT_STOP_, SSRT_STOP-CHANGE_) in groups of healthy participants ranging from 8 to 24 individuals (Boecker et al., 2013). We thus targeted 24 participants as the upper range of the previously reported experiments. The SSRT_STOP_ vs. SSRT_STOP-CHANGE_ difference mean effect size was 0.65 across the reviewed experiments. Including this value together with a pre-defined alpha level of .05 and a 90% statistical power, a sample size calculation with G*Power (Faul et al., 2009) confirmed that our sample was sufficient to investigate the anticipated effect.

#### Experimental task

The task design followed recent guidelines for stopping tasks procedures (Verbruggen et al., 2019). The participants performed a randomized sequence of STOP-task blocks and STOP-CHANGE-task blocks involving foot responses within a single session. The behavioral task was presented on a Linux desktop computer based on Matlab 2017a (The MathWorks) custom scripts, including Psychophysics Toolbox functions (Brainard, 1997). The task design is depicted in **Figure 1**. The stimuli were displayed centrally on a black background screen. The main GO task was a two-choice reaction-time task. In each trial, after a fixation cross was displayed for a random duration ranging from 400 to 700 ms, a white arrow pointing to the left or the right direction was displayed as the GO stimulus. Participants were instructed to respond quickly and accurately to the GO stimulus by pushing the corresponding left or right foot pedal on a Kinesis Savant Elite 2 response device. In a pseudorandomly chosen 1/3 of the trials, the white arrow (GO) turned into a STOP or STOP-CHANGE signal after a variable delay (see below). The STOP signal was a blue rectangle instructing participants to try to stop the initial GO response. The STOP-CHANGE signal was a blue arrow pointing in the opposite direction instructing participants to try to stop the initial GO response and, instead, push the opposite foot pedal. Therefore, the STOP-CHANGE instruction included both the requirement to stop the initial response and an additional demand to replace it with another response. The STOP/ STOP-CHANGE signal-delay (SSD) between the onset of the GO stimulus and the STOP/ STOP-CHANGE signal display, initially set to 200 ms, was dynamically adjusted in 50 ms increments to achieve a probability of responding to the initial GO stimulus p(resp|signal) of ∼.50 in signal trials. If the participant completed the initial GO response after a STOP/ STOP-CHANGE signal presentation, the trial was considered unsuccessful and the SSD was shortened in the next signal trial (US: Unsuccessful STOP; UC: Unsuccessful STOP-CHANGE). Otherwise, if the initial GO response was canceled, the trial was considered successful and the SSD was lengthened in the next signal trial (SS: Successful STOP; SC: Successful STOP-CHANGE). The incremental SSD adjustment was specific to the Block Type (stopping or changing) and to the responding foot (left or right). If the reaction time to the GO stimulus or STOP-CHANGE signal was above 1000 ms, the feedback ‘Too slow!’ was displayed on the screen for a 1300 ms duration. Overall, the duration between two subsequent GO stimuli was 3600 ms.

**Figure 1.**
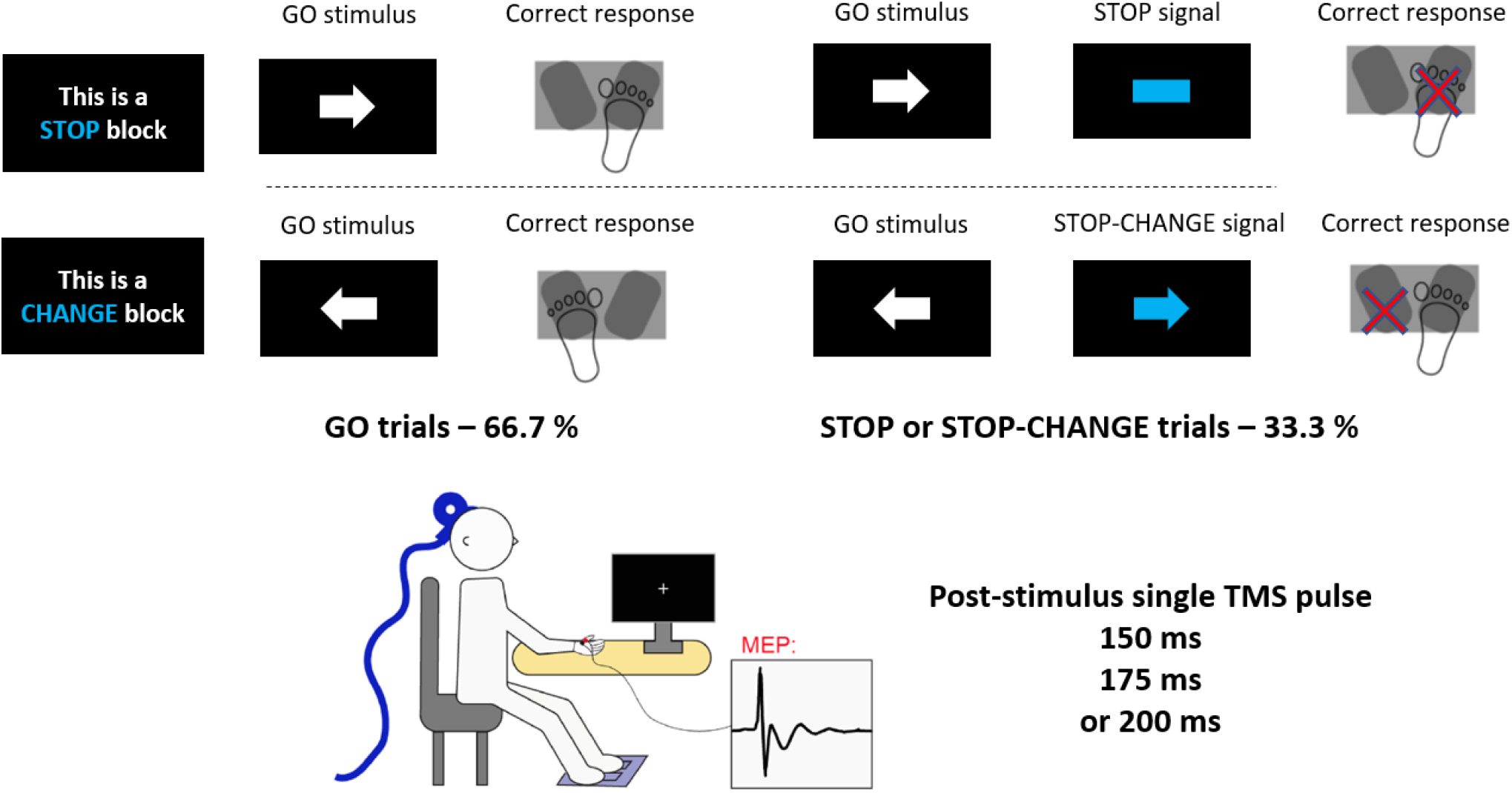
Task diagram of Experiment 1. Top, Time course of the behavioral task. Following presentation of the GO stimulus, the arrow changed to blue on a minority of trials (33.3%). These signaled to the participant to try to stop their initial response (STOP signal) or to stop the initial response for changing it with the newly instructed response (STOP-CHANGE signal). Bottom, the setup which involved recording MEPs from the right hand while the feet were used to respond.

Participants performed 12 blocks of 90 trials (60 GO trials + 30 STOP/ STOP-CHANGE signal trials). Instructions for the upcoming block (either STOP-task or STOP-CHANGE-task block) were displayed before each block initiation. Blocks started with a sequence of three consecutive GO trials and were concluded by written and verbal feedback including the block average GO-trials reaction time (GO-RT) and p(resp|signal). To maintain a p(resp|signal) close to .50 and between .40 and .60, the participants were encouraged either to speed up their GO-RT or to increase their successful stopping/changing rate. A minimum of two practice blocks (stopping and changing), 30 trials each (10 signal trials), were performed before the actual experiment. This practice was repeated if the p(resp|signal) was outside the .40 to .60 range. White and blue color correspondence to GO stimuli and STOP/ STOP-CHANGE signals was randomized across participants.

#### TMS/MEP procedures

CSE was assessed based on an EMG measure of the motor-evoked potential (MEP) induced by a single TMS pulse. As we use the MEP to investigate a non-selective inhibition process, it was quantified from the right index finger as a task-irrelevant effector (Badry et al., 2009; Majid et al., 2012; Wessel et al., 2013).

EMG was measured using a bipolar belly-tendon montage from the right first dorsal interosseous muscle using adhesive electrodes (H124SG, Covidien) and a ground electrode placed over the distal end of the ulna bone. EMG electrodes were passed through a Grass P511 amplifier (Grass Products) and sampled using a CED Micro 1401-3 sampler (Cambridge Electronic Design) and CED Signal software. EMG was sampled at a rate of 1000 Hz with online 30 Hz high-pass, 1000 Hz low-pass, and 60 Hz notch filters. EMG sweeps were triggered 90 ms before each TMS pulse, and EMG was recorded for 1000 ms. TMS was delivered via a MagStim 200-2 system (MagStim) with a 70 mm figure-of-eight coil. Before the experimental task, hotspotting was used to identify the correct location and intensity for each participant. The coil was initially located five cm left of and two cm anterior to the vertex. Location and stimulation intensity were incrementally adjusted to identify the participant’s resting motor threshold. Resting motor threshold was the minimum intensity needed to produce MEPs > 0.1 mV in 5 of 10 consecutive probes (Rossini et al., 2015). For the main experiment, the stimulus intensity was increased to 115% of resting motor threshold. The mean experimental intensity was 62% (SD = 7.6%) of maximum stimulator output.

Participants completed 1080 trials. In 1/3 of the GO trials (n = 240), the TMS pulse was delivered 50 ms prior to the GO stimulus display to measure an active baseline MEP. Another 1/3 of the GO trials were TMS-pulse free (n = 240). In the last 1/3 of GO trials (n = 240), as well as every STOP (n = 180) and STOP-CHANGE trial (n = 180), a TMS pulse was delivered randomly at one of the three following latencies: 150 ms, 175 ms, 200 ms (resulting in n = 60 trials per signal type for a given latency). The TMS-pulse latencies were chosen based on previous work establishing MEP suppression as a marker of non-selective inhibition through the cortico-subcortical hyperdirect pathway (Tatz et al., 2021; Wessel et al., 2022).

#### Data analyses

GO_STOP_-RT and GO_CHANGE_-RT were computed as the mean GO-RT from STOP and STOP-CHANGE blocks, respectively. US-RT and UC-RT were computed as the mean RT of unsuccessful STOP trials and unsuccessful STOP-CHANGE trials (STOP-CHANGE trials in which the original response was executed), respectively. GO2-RT was also computed as the mean RT to the STOP-CHANGE signal when changing was successful. Participants’ behavioral inhibition latencies (STOP/ STOP-CHANGE-signal reaction time), SSRT_STOP_ and SSRT_CHANGE_, were estimated using the integration method with replacement of GO omissions (Verbruggen and Logan, 2009; Verbruggen et al., 2019). This consisted of subtracting the mean SSD from the *n*^th^ RT, where *n* equals the number of GO-RTs multiplied by the overall p(resp|signal). Blocks with a p(resp|signal) outside the .30 to .70 range were excluded. Overall, this was the case for three STOP blocks (across three participants) and three STOP-CHANGE blocks (across two participants). Trials with a GO-RT or GO2-RT shorter than 150 ms or longer than 1000 ms were also removed. The average number of removed trials per participant was 2.3 (SD = 6.3) for stopping and 2.9 (SD = 6.8) for changing.

MEP amplitude was extracted semiautomatically from the EMG trace using ezTMS, a freely available TMS preprocessing tool developed for use in MATLAB (Hynd et al., 2021; Tatz et al., 2021). MEP amplitude was defined as the difference between the maximum and minimum amplitude 10-50 ms after the TMS pulse. In addition to automatically determining MEP amplitude, each trial was visually inspected for accuracy without knowledge of the specific trial type. Trials in which the MEP amplitude was < 0.01 mV or the root mean square (RMS) of the EMG trace was > 0.01 mV were excluded from the MEP analyses. The first 80 ms of the EMG sweep were used to calculate the RMS power. Mean MEP amplitudes were computed for each condition for each participant. These values were then normalized by dividing them by the mean active baseline MEPs (pre-GO TMS pulse). Across participants, an average number of 29 GO MEPs per TMS-pulse latency and per Block Type (STOP and STOP-CHANGE) were included in further analysis. In signal trials, there were, on average, 52 resulting MEPs per TMS-pulse latency and Block Type.

#### Statistical analysis

To investigate differences between stopping- and changing-conditions measurements, we used two-tailed paired *t*-tests. When assessing the effect of a task-design manipulation or investigating an a priori hypothesized effect, one-tailed tests were applied (specified in the results section). Pearson correlations were also performed across participants to assess the relationship between behavioral and neurophysiological measurements. When multiple tests were used, a false-discovery rate (FDR) correction (Benjamini et al., 2006) was applied to *p*-values to account for type I error inflation. For significant differences, a 95% confidence interval and Cohen’s D effect size were reported. For non-significant results, we reported the Bayes factor evidence for the null hypothesis (BF_01_) (Krekelberg, 2023).

#### Data availability

The data files, as well as the scripts for running the task and analyzing the data can be found on the Open Science Framework at [*URL to be inserted after acceptance*].

### Results

#### Behavior

The main behavioral and CSE results for Experiment 1 are presented in **Figure 2**. Values for p(resp|signal) did not differ between STOP and STOP-CHANGE tasks (paired *t*-test, *t_(23)_* = 1.01, *p* = .32, BF_01_ = 2.94), and were in both cases not significantly different from .50 (STOP: *t_(23)_* = 0.16, *p* = .87, BF_01_ = 4.60, STOP-CHANGE: *t_(23)_* = 0.85, *p* = .40, BF_01_ = 3.36). In addition, US-RT and UC-RT were significantly shorter than GO_STOP_-RT (one-tail paired *t*-test, *t_(23)_* = 6.74, *p* < .001, CI_95_[40 ms, +∞], *Cohen’s d* = 0.75) and GO_STOP-CHANGE_ RT (*t_(23)_* = 7.81, *p* < .001, CI_95_[39 ms, +∞], *d* = 0.77), respectively. These manipulation-checks verified that the assumptions of the race model held (Verbruggen et al., 2019) and permitted reliable SSRT_STOP_ and SSRT_STOP-CHANGE_ estimations.

**Figure 2.**
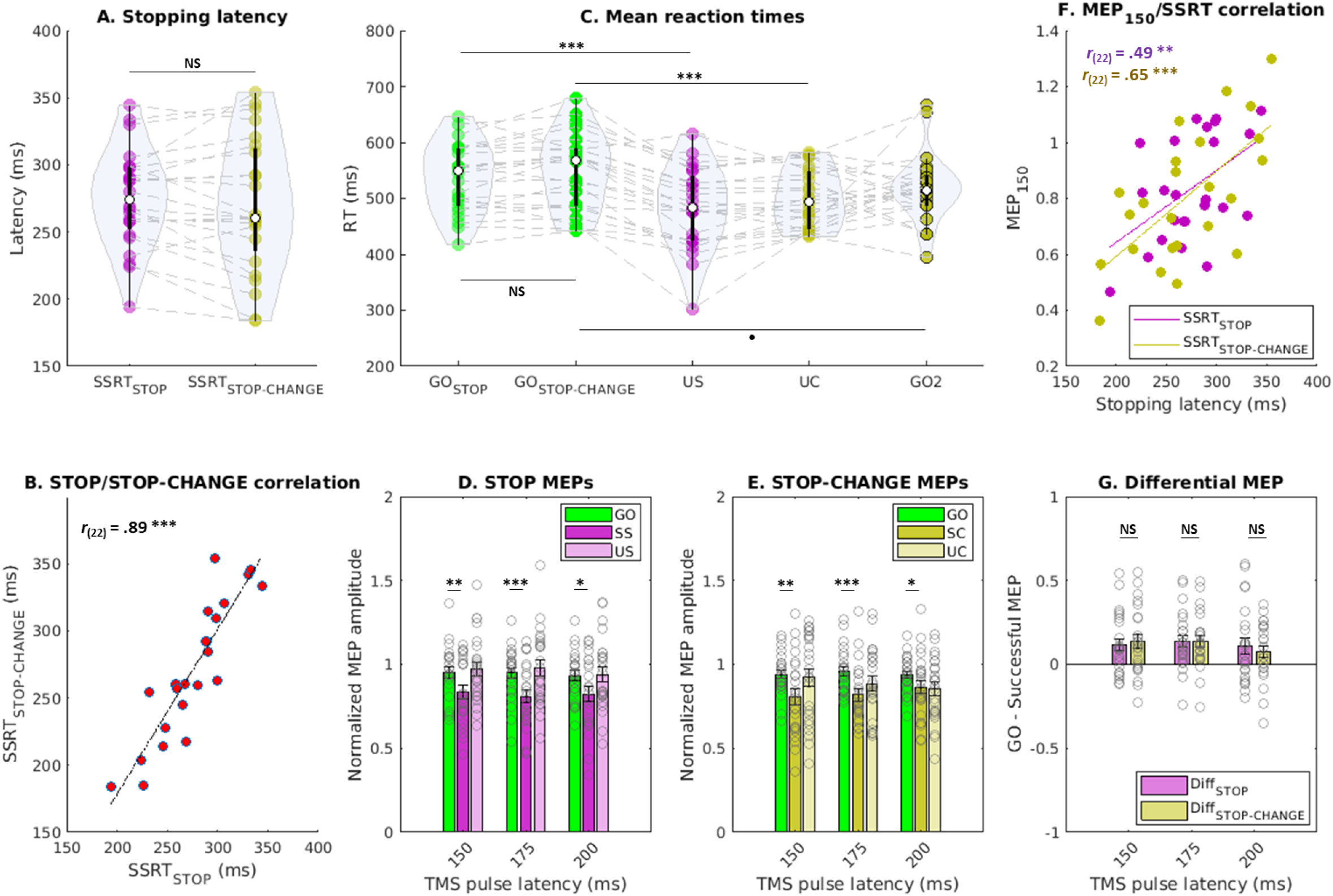
Behavior and CSE results of Experiment 1. Panel A and B, behavioral response stopping latencies (SSRTs) estimated for action-stopping and -changing, and their correlation across participants. Panel C, individuals’ mean RT in GO, unsuccessful STOP/STOP-CHANGE (US, UC), and successfully changed responses (GO2). Panel D and E, MEPs amplitudes in STOP- and STOP-CHANGE-task blocks, respectively. Panel F, MEPs/behavior correlations. Panel G, GO vs. Successful signal-trials MEPs contrast in stopping and changing. ^●^ *p* = .05, \**p_FDR_* < .05, ** *p_FDR_* < .01, *** *p_FDR_* < .001.

The average participants’ GO trials accuracy was .98 (SD = .03) in STOP blocks and .97 (SD = .02) in CHANGE blocks. The difference between GO_STOP_-RT (M = 537 ms, SD = 61 ms) and GO_STOP-CHANGE_-RT (M = 550 ms, SD = 67 ms) was not significant (*t_(23)_* = 2.00, *p* = .057, BF_01_ = 0.85) and the two values were highly correlated across participants (*Pearson’s r_(22)_* = .89, *p* < .001, CI_95_[.76, .95]). These results indicated that action initiation was performed similarly across Block Type, and that higher proactive slowing of the initial GO response in change-vs. stop-task blocks was minimal, if any.

The behavioral inhibition latencies, SSRT_STOP_ (M = 275 ms, SD = 36 ms) and SSRT_STOP-CHANGE_ (M= 270 ms, SD = 50 ms), did not differ significantly (*t_(23)_* = 0.98, *p* = .33, BF_01_ = 3.01) and were highly correlated (*r_(22)_*= .89, *p* < .001, CI_95_[.77, .95]), suggesting commonalities in the response cancellation required in STOP and STOP-CHANGE trials. In STOP-CHANGE blocks, the GO2-RT (M = 518 ms, SD = 60 ms) was tendentially (no statistical significance) shorter than GO_CHANGE_-RT (*t_(22)_* = 2.03, *p* = .05, CI_95_[-1 ms, 64 ms], *d* = 0.49), suggesting that response changing might be faster than its primary initiation. Overall, these results suggest a broad commonality of processing across STOP and STOP-CHANGE blocks.

#### Non-selective CSE suppression

First, we verified that the baseline MEP amplitude did not differ between STOP and STOP-CHANGE blocks, which it did not (*t_(23)_* = 1.30, *p* = .20, BF_01_ = 2.20). In addition, the GO MEP amplitude did not differ across Block Type and latencies (all uncorrected *p* values > .68).

Second, we assessed the significance of the expected non-selective MEP reduction in STOP blocks to replicate previous findings of global CSE suppression in human-action stopping (Badry et al., 2009; Wessel et al., 2022). MEP amplitude was significantly lower in SS than GO trials at all three latencies (one-tail paired *t*-test with FDR correction, 150 ms: *t_(23)_* = 3.16, *p_FDR_* = .003, CI_95_[0.05, +∞], *d* = 0.63; 175 ms: *t_(23)_* = 3.98, *p_FDR_*< .001, CI_95_[0.08, +∞], *d* = 0.73; 200 ms: *t_(23)_*= 2.40, *p_FDR_* = .01, CI_95_[0.03, +∞], *d* = 0.58).

Third, the main hypothesis of Experiment 1 was that non-selective CSE suppression would also occur in STOP-CHANGE trials. The hypothesis was verified, as SC trials MEPs were reduced as compared to GO_CHANGE_ trials (one-tail paired *t*-test, 150 ms: *t_(23)_* = 3.28, *p_FDR_* = .002, CI_95_[0.06, +∞], *d* = 0.68; 175 ms: *t_(23)_*= 4.02, *p_FDR_* < .001, CI_95_[0.08, +∞], *d* = 0.81; 200 ms: *t_(23)_* = 2.08, *p_FDR_* = .02, CI_95_[0.01, +∞], *d* = 0.46).

Fourth, to compare the level of non-selective CSE suppression engaged in successful action-stopping and -changing, we compare SS vs. SC trials MEPs. This contrast showed no difference across the three TMS pulse latencies (all uncorrected *p* values > .26). In addition, SS and SC MEPs values were significantly correlated across participants at the three TMS latencies (150 ms: *r_(22)_* = .75, *p_FDR_* < .001, CI_95_[.50, .89]; 175 ms: *r_(22)_* = .76, *p_FDR_* < .001, CI_95_[.51, .89]; 200 ms: *r_(22)_*= .62, *p_FDR_* = .001, CI_95_[.29, .82]). We further computed for each Block Type the difference (ΔMEP) between successful STOP/CHANGE and GO trials MEPs. This contrast indicated a similar level of global CSE suppression between the two Block Type as ΔMEP_STOP_ and ΔMEP_STOP-CHANGE_ did not differ (paired *t*-test, 150 ms: *t_(23)_* = 0.53, *p* = .60, BF_01_ = 4.09; 175 ms: *t_(23)_* = 0.03, *p_FDR_* = .98, BF_01_ = 4.66; 200 ms: *t_(23)_*= 0.85, *p* = .40, BF_01_ = 3.36). To the best of our knowledge, this experiment is the first to show a non-selective CSE suppression evoked by action-changing. Bayes factor indicated moderate evidence for similar suppression levels between stopping and changing.

Fifth, we assessed the behavioral relevance of non-selective CSE suppression at the earliest latency (150 ms). The MEP amplitude in SS and SC trials was significantly correlated to SSRT_STOP_ (*r_(22)_* = .49, *p* = .01, CI_95_[.11, .75]) and SSRT_STOP-CHANGE_ (*r_(22)_* = .65, *p* < .001, CI_95_[.33, .83]), respectively, reflecting that larger CSE suppression was associated with faster response cancelation in both stopping and changing tasks. Although the non-selective CSE suppression has been considered a signature of broad motoric inhibition in previous action-stopping studies (Duque et al., 2017; Wessel and Aron, 2017), our results demonstrate for the first time that this early marker is directly associated with the behavioral inhibition latency (SSRT). We further demonstrate that this relation is also valid in action-changing.

In summary, these results suggest that both infrequent STOP and STOP-CHANGE signals feature non-selective CSE suppression (‘pause’) contributing to the fast revision of the action.

## Experiment 2 & 3

### Method

#### Hypotheses

Experiment 1 showed that non-selective CSE suppression reflects a global ‘pause’ stage that is commonly triggered in action-stopping and -changing. Experiment 2 and 3 further aimed at disentangling additional proposed neurophysiological signatures of inhibitory control. Based on ‘pause-then-cancel’ (PTC) predictions, we anticipated that 1) frontocentral EEG β-bursts dynamics and partial EMG responses act as neurophysiological correlates of the early ‘pause’ stage (Tatz et al., 2021, 2023; Wadsley et al., 2023a) across both situations (stopping and changing), and 2) the amplitude of ‘cancel’ inhibition-stage signatures, including frontocentral P3 ERP and θ power (Diesburg and Wessel, 2021; Messel et al., 2021), scale with the required action-revision (see below).

For the EEG and EMG investigations, we used a task in which STOP and CHANGE trials were intermixed within task blocks. Indeed, previous works have suggested that the block design modulates the SSRT_STOP_ vs. SSRT_STOP-CHANGE_ difference (Boecker et al., 2013). After showing similar SSRT_STOP_ and SSRT_STOP-CHANGE_ in a between-block design (Experiment 1), Experiment 2 and 3 compared these measures in a within-block design. Furthermore, beyond the action-stopping vs. -changing comparison, we investigated whether EEG purported signatures of inhibitory control would be affected by modulating the amount of action-change. Therefore, we included two types of STOP-CHANGE signals. In STOP-CHANGE-side (CS) trials, participants had to change one response dimension, similar to Experiment 1 (see below). In STOP-CHANGE-side-and-finger (CSF) trials, participants had to change two dimensions of the response (see below). In Experiment 2, the 1/3 of trials in which a signal was presented were divided by the three Signal Types: 1/9 STOP, 1/9 CS, 1/9 CSF. Then, to ensure that our results were not led by the higher probability of CHANGE trials (2/9 for CS + CSF count) compared to STOP trials (1/9), we aimed to replicate our findings in a separate experiment. That experiment used the exact same task design in Experiment 3, but included a modification of these probabilities. The 1/3 of signal trials were divided into half STOP trials (1/6) and half STOP-CHANGE trials (1/12 CS and 1/12 CSF). Empirically, these probabilities did not influence any of the results in a significant fashion. We will therefore present data from the combined sample (N = 52) to achieve the highest amount of statistical power and fidelity.

#### Participants

30 healthy individuals (16 females, mean age 19.4 years, SD = 1.2) participated in Experiment 2 and twenty-two (16 females, mean age 21.6 years, SD = 4.3) in Experiment 3, respectively, in exchange for course credit or an hourly payment of $15. All were right-handed and had normal or corrected-to-normal vision. None of the participants reported a history of psychiatric or neurological disorders. The experiments were performed with the informed consent of all participants, and procedures were approved by the local Institutional Review Board (IRB #201612707) and conducted in accordance with the Declaration of Helsinki.

#### Experimental task

The behavioral task was presented on a Linux desktop computer using on Matlab 2017b (The MathWorks) and the Psychophysics Toolbox (Brainard, 1997). The task design is depicted in **Figure 3**.

**Figure 3.**
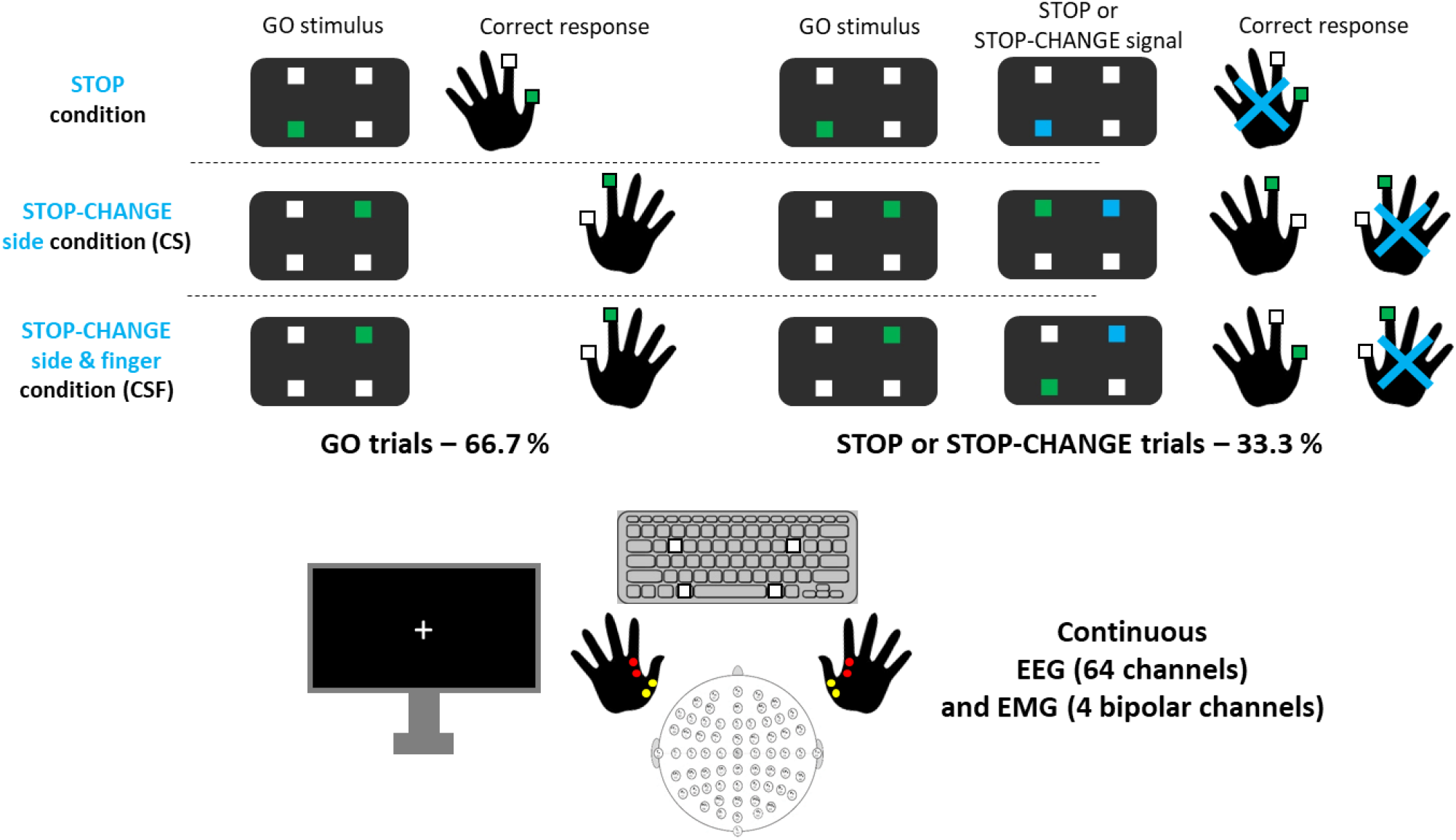
Task diagram of Experiment 2 & 3. Time course of the behavioral task. Following the presentation of the green GO target, the target changed to blue in a minority of trials (33.3%). These signaled the participant to try to stop their initial response (STOP signal) or stop the initial response for changing it with the newly instructed response: another green target (STOP-CHANGE-side or STOP-CHANGE-side-and-finger signal). Setup additionally involved a scalp EEG recording and EMG recordings of first-dorsal interosseous muscles for index fingers (red) and abductor pollicis brevis muscles for thumbs (yellow).

The main GO task was a 4-choice reaction-time task. In each trial, after four white squares were presented for a random duration ranging from 500 to 750 ms, one of the squares turned green, representing the GO stimulus. Participants were instructed to respond quickly and accurately to the GO stimulus by pressing the corresponding key on the response device, a keyboard with 4 keys colored in white to match the four potential GO stimuli spatially. Responses were made with the right/left index finger or thumb. In a randomly chosen 1/3 of the trials, the initial green square (GO) turned into a STOP or STOP-CHANGE signal after the SSD. If the green square turned blue, this was a STOP signal, instructing participants to try to cancel the GO response. If the initial green square turned blue while another white square turned green at the same time, this was a STOP-CHANGE signal, instructing participant to try to cancel the initial GO response and, instead, press the key corresponding to the new green stimulus. The STOP-CHANGE signal always engaged a switch in the response side (e.g., left to right), but could either engage an additional finger switch (left index finger to right thumb) or no finger switch (left index finger to right index finger), thus implying to modify two vs. one dimensions of the response. Therefore, three Signal Type could have been displayed after the initial GO stimulus: STOP signal, CS signal (STOP-CHANGE-side), CSF signal (STOP-CHANGE-side-and-finger). Specific to each Signal Type and each finger, the SSD incremental adjustment was similar to Experiment 1 to achieve a probability of responding p(resp|signal) of ∼.50 in signal trials. Trials were categorized as GO trials, SS/SCS/SCSF trials (successful stopping and changing trials), or US/UCS/UCSF (unsuccessful stopping and changing trials). Erroneous responses (i.e., pressing another key than the cued GO or CHANGE responses) were excluded (< 1% of the data). If the reaction time to the GO stimulus or STOP-CHANGE signal was above 1000 ms, the feedback ‘Too slow!’ was displayed on the screen for a 500 ms duration. Overall, the duration between two subsequent GO stimuli was 3000 ms. The participants performed 1008 trials through 14 blocks of 72 trials (48 GO trials + 24 signal trials). Each block started with three consecutive GO trials, and written and verbal feedback was provided after each block as described in Experiment 1. Before the actual task, four practice blocks of 24 trials each were performed: one GO-only block, one GO+STOP block, one GO+STOP-CHANGE block, one intermixed block. This practice was repeated if the p(resp|signal) was outside the .40 to .60 range. Green and blue color correspondence to GO stimuli and STOP/ STOP-CHANGE signals was randomized across participants.

#### EEG and EMG recording

EEG was acquired using a 64-channel active electrode cap connected to an ActiChamp amplifier (Brain Products). The ground and reference electrodes were placed at AFz and Pz, respectively. EMG was recorded from the first-dorsal interosseous muscles for index fingers and abductor pollicis brevis muscles for thumbs. For each, two electrodes were placed on the belly and tendon of the muscle. Ground electrodes were placed on the distal end of each hand’s ulna (for indexes) and radius (for thumbs). The EMG electrodes were connected to the EEG system using two auxiliary channels (via BIP2AUX adaptor cables). Thus, there were 68 total recording channels. All channels were sampled at a rate of 2500 Hz. To minimize task-unrelated EMG activity, the participants were instructed only to move their fingers when necessary and to keep both hands otherwise relaxed and pronated on the desk.

#### EEG and EMG preprocessing

Custom MATLAB scripts, including EEGLAB (Delorme and Makeig, 2004) toolbox functions, were used to preprocess the EEG and EMG data. Both EEG and EMG data were bandpass filtered (EEG: 0.1-50 Hz; EMG: 20-250 Hz) and downsampled to a rate of 500 Hz. The continuous EEG data were visually inspected to identify non-stereotypical artifacts. Segments with non-stereotypical EEG artifacts were removed from the EEG and EMG data. EEG and EMG data were epoched relative to the onsets of GO stimuli (-500 to 2000 ms). Missed and incorrect GO trials were excluded from all analyses. The EEG data were re-referenced to the common average. Independent component analysis (Infomax ICA; Bell & Sejnowski, 1995) was applied to the continuous EEG data (concatenation of the EEG epochs) to identify neural components contributing to the observed scalp data. Using the ICLABEL classifier (Pion-Tonachini et al., 2019), the components associated with an above .90 probability to be artifactual were removed from the EEG data structure, thus removing their contributions to the observed EEG. Rejection was systematically checked by visual inspection of component properties (time series, spectra, topography) according to ICLABEL guidelines. The EMG data were then converted to RMS power using a sliding window of 10 sample points and baseline-corrected by dividing the entire epoch by the mean of the 200 ms prestimulus EMG. The resulting data were then standardized for each finger across all types of trials and sample points using the z-transform.

#### Behavioral analysis

Similar to Experiment 1, we computed mean participants’ reaction times: GO-RT (GO trials), US-RT/UCS-RT/UCSF-RT (Unsuccessful STOP/STOP-CHANGE trials), GO2_CS_-RT/GO2_CSF_-RT (new response for successful STOP-CHANGE trials). The behavioral inhibition latency was also estimated according to Signal Type: SSRT_STOP_/SSRT_CS_/SSRT_CSF_.

#### EMG data analysis

As a manipulation check, we examined whether the EMG data accurately captured the participant’s motor responses on GO trials. Single-trial GO-RTs were correlated to EMG peak latencies within each EMG channel (*Pearson’s r*). The EMG peak was computed as the maximal value within the RT window. For those correlations, trials in which *Cook’s D* exceeded 4/N, with N the number of channel’s data points, were deemed statistical outliers and excluded from further analysis. Then, channels with a *r* value lower than .5 were also excluded, due to potential disconnections of the electrodes during the experiment. Participants with more than one rejected channel were excluded from the analysis.

For each condition, the trial-to-trial EMG peak was identified, as explained above. This procedure was applied for trials with a behavioral response (GO, US, UCS and UCSF trials). For trials with no response to the primary target (SS, SCS and SCSF trials), we aligned with previous studies to identify partial EMG (prEMG) responses (Jana et al., 2020; Tatz et al., 2021; Raud et al., 2022). Within a window corresponding to the maximal RT of each participant, an EMG burst was identified in a trial if any data point exceeded a threshold of 1.2 (in standard deviation units, taking into account all trials) (Raud et al., 2020, 2022). The peak latency was defined as the time-point of the highest amplitude in a trial. Lastly, peak latencies of the EMG were recalculated relative to the stop signal onset by subtracting the trial SSD.

#### EEG data analysis

Previous studies have shown that β-burst dynamics convey inhibitory commands along cortico-subcortical networks, from STN to M1 (Cagnan et al., 2019; Diesburg et al., 2021), and from frontocentral sites to sensorimotor sites (Wessel, 2020; Choo et al., 2022). We thus computed frontocentral β-bursts as an index of early cortical inhibitory mechanisms (Hannah et al., 2020; Jana et al., 2020; Tatz et al., 2023). The FCz electrode’s data were convolved with a complex Morlet wavelet of the form:

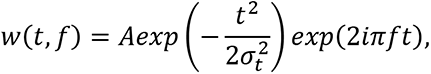

*with* 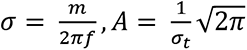, and *m* (cycles) ranging from five to nine across the 15 evenly spaced frequencies spanning the β-band (15–29Hz). Time-frequency power estimates were extracted by calculating the squared magnitude of the complex wavelet-convolved data. Individual β-bursts were identified in the trial-by-trial β-band time-frequency power matrix based on power exceeding a cutoff of 2× the median power of the entire time-frequency power matrix for that electrode. Individual’s β-burst rates were computed for each condition based on a 50ms-length sliding window (Choo et al., 2022; Tatz et al., 2023). Time points wise, the number of bursts across trials was summed within the window and divided by the number of trials. β-burst volume was computed for each burst through numerical integration of the suprathreshold datapoints (Enz et al., 2021). A similar sliding-window averaging resulted in a single vector reflecting the β-burst volume time-dynamic for each condition and each participant. These β-burst rate and volume time series were then relocked for each condition to the onset of the STOP/STOP-CHANGE signal or the match-GO latency (GO stimulus onset + average participant’s SSD). Individuals’ vectors were normalized by dividing the values by the average of the -200 to 0 ms baseline period. Then, we computed individuals’ average normalized β-burst rate and volume within each participant’s SSRT window.

Second, EEG correlates of action-stopping are classically observed at later latencies with stop-signal-related frontocentral P3 wave and θ power increase (Huster et al., 2013; Wessel and Aron, 2015; Waller et al., 2019; Dykstra et al., 2020; Messel et al., 2021). According to extant theoretical proposals, both are thought to index a later selective stage of the inhibitory control that specifies the need to cancel the response (Waller et al., 2019; Diesburg and Wessel, 2021; Tatz et al., 2021). Individuals’ frontocentral ERPs were obtained for each condition by averaging FCz site EEG data across trials. To compute θ power, we band-pass filtered FCz data in the 4-8 Hz frequency range. The filtered time series were then translated into complex space using the Hilbert transform (as implemented in the MATLAB *hilbert* function). θ power was then extracted by computing the square of the absolute of the complex output of the Hilbert transform. Individuals’ ERPs and θ power were re-epoched into a -200 to 800 ms surrounding the STOP/ STOP-CHANGE signal onset (or match GO), and we subtracted a baseline defined as the average signal in the -200 to 0 ms window. P3 and θ amplitudes were defined as the maximal value of the ERP and Power time series. Following Wessel and Aron (2015), the P3 onset was defined as the earliest time point at which a statistically significant deviation between STOP/ STOP-CHANGE and matched GO trials ERPs could be detected in a given participant. For this, *t*-tests were performed across single-trial EEG. Working back over data samples from the P3 peak, the earliest sample point at which the STOP/CHANGE ERP significantly exceeded the GO ERP was selected as the participant’s P3 onset. Individuals’ ERP peaks and onsets were successfully computed in all task conditions for most of the participants. When an individual’s value was unsuccessfully captured for a given condition, this participant was removed from the statistical comparisons involving this condition. Therefore, degrees of freedom ranged from 40 to 50 for the statistical comparisons we performed.

Third, we completed our comparative analysis of the neurophysiological dynamic of action -stopping vs. -changing by investigating at which time point a multivariate pattern analysis (MVPA) classifier applied to the whole-scalp EEG would be able to distinguish between the conditions of interest above chance. MVPA analyses were performed using the ADAM toolbox (Fahrenfort et al., 2018) on the preprocessed 64 channels ‘data. We used leave-one-out cross-validation in which a Linear Discriminant Analysis classifier was trained on all trials, but one, and tested on the remaining trial that was not part of the training set. This validation was implemented in a sample-point-wise fashion, in which training and testing were done at each sample point following the STOP/CS/CSF signal. Classifier performance was measured as the area under the receiver operating characteristics curve (AUC) (Wickens, 2002), which quantified the total AUC when the cumulative true positive rate (probability of correct classification) was plotted against the false positive rate (probability of incorrect classification) for each decoding problem between the conditions of interest. Time series of AUC values were obtained for each pair of conditions and each participant. To identify time points during which classification performance was significantly above chance, we compared each AUC value against chance-level performance (0.5) using one-sample *t*-tests, which was corrected for multiple comparisons using cluster-based permutation testing (10,000 iterations, cluster-*p* value = .00001, individual sample point *a* = .0001) (Maris and Oostenveld, 2007).

### Results

The results of experiments 2 and 3 converged toward a highly similar pattern. Therefore, the analyses presented below were performed on the two experiments’ data altogether (N = 52) to increase statistical power.

#### Behavior

The main behavioral results of Experiment 2 and 3 are depicted in **Figure 4**. One participant was rejected because of an outlier SSRT_STOP_ value (< M ± 3 SD). Final sample for the behavioral analysis was N = 51. The average participants’ GO trials accuracy was .98 (SD = .02). In STOP trials, p(resp|signal) (M = .46, SD = .05) was significantly lower than CS trials (M = .51, SD = .02; paired *t*-test with FDR correction, *t_(50)_* = 7.01, *p_FDR_* < .001, CI_95_[.03, .06], *Cohen’s d* = 1.01) and CSF trials (M = .51, SD = .02; *t_(50)_* = 6.62, *p_FDR_* < .001, CI_95_[.03, .06], *d* = 0.99). CS and CSF conditions did not differ (*t_(50)_* = 0.32, *p_FDR_* = .75, BF_01_ = 6.26). These results indicate a higher successful cancellation rate of the initial-response in action-changing compared to -stopping (despite adaptive tracking being used for both).

**Figure 4.**
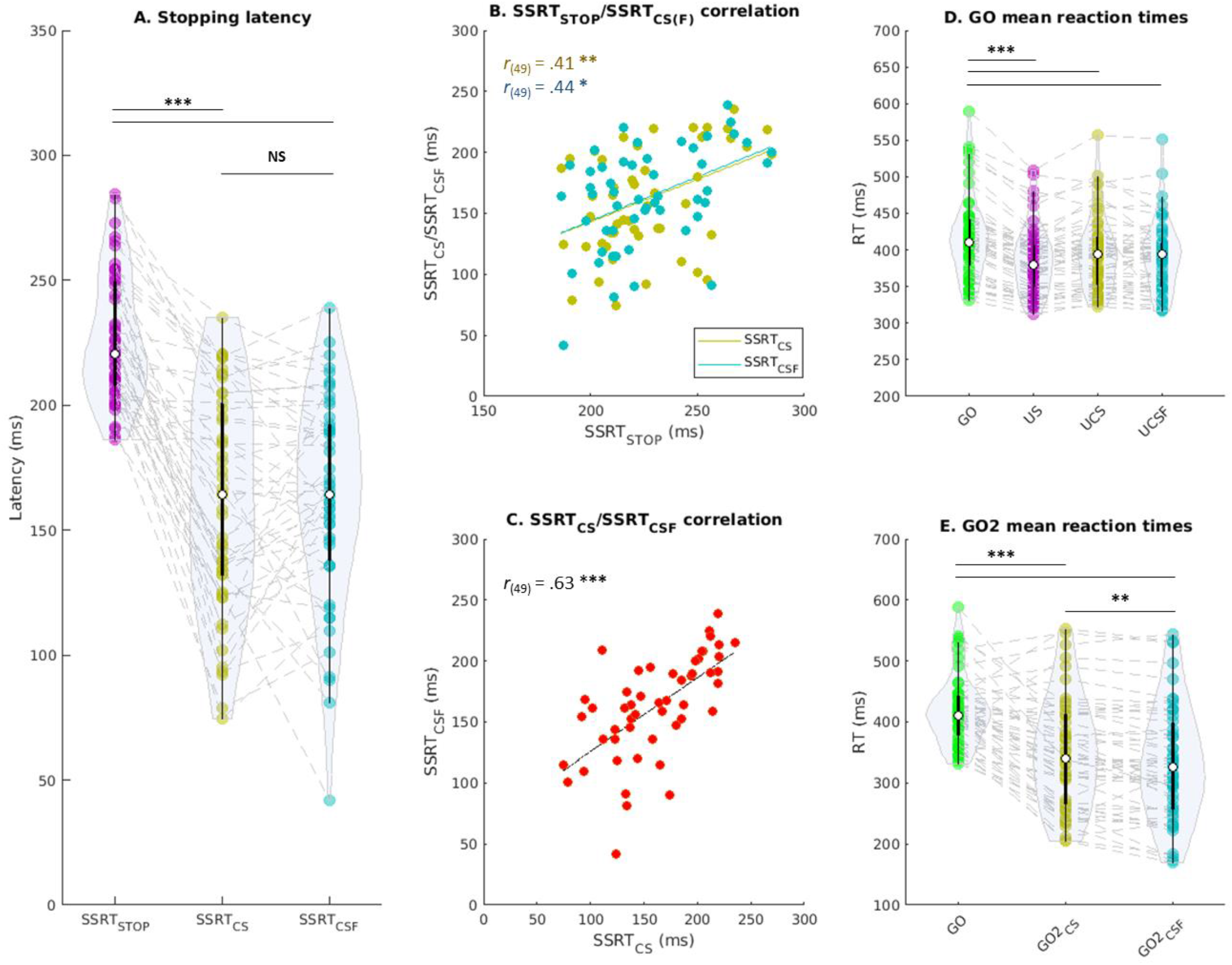
Behavior results of Experiment 2 & 3. Panel A, behavioral initial-response inhibition latencies (SSRTs) estimated for action-stopping and -changing. Panel B, correlations between STOP and STOP-CHANGE SSRT estimates across participants. Panel C, correlation between Change-side and Change-side-and-Finger SSRT estimates across participants. Panel D, individuals’ mean RT in GO and unsuccessful STOP/STOP-CHANGE (US, UCS, UCSF) responses. Panel E, individuals’ mean RT in successfully changed responses (GO2_CS_, GO2_CSF_). \**p_FDR_* < .05, ** *p_FDR_* < .01, *** *p_FDR_* < .001.

Verifying race model assumptions (Verbruggen et al., 2019), GO-RT (M = 418 ms, SD = 57 ms) was significantly longer than US-RT (M = 386 ms, SD = 46 ms; one-tail paired *t*-test, *t_(50)_* = 9.87, *p_FDR_* < .001, CI_95_[27 ms, +∞], *d* = 0.60), UCS-RT (M = 396 ms, SD = 51 ms; *t_(50)_* = 6.25, *p_FDR_*< .001, CI_95_[16 ms,, +∞], *d* = 0.41), and UCSF-RT (M = 391 ms, SD = 48 ms; *t_(50)_* = 8.22, *p_FDR_* < .001, CI_95_[21 ms,, +∞], *d* = 0.50). These manipulation-checks verified that the assumptions of the race model held and permitted reliable SSRT_STOP_ and SSRT_CHANGE_ estimations.

Suggesting a faster initial-response cancellation in action-changing than -stopping, SSRT_STOP_ (M = 226 ms, SD = 25 ms) was significantly longer than SSRT_CS_ (M= 161 ms, SD = 43 ms; *t_(50)_* = 11.60, *p_FDR_* < .001, CI_95_[54 ms, 76 ms], *d* = 1.35) and SSRT_CSF_ (M= 163 ms, SD = 41 ms; *t_(50)_* = 12.00, *p_FDR_* < .001, CI_95_[53 ms, 74 ms], *d* = 1.36) while SSRT_CS_ and SSRT_CSF_ did not differ (*t_(50)_* = 0.29, *p_FDR_* = .77, BF_01_ = 8.16). A moderate correlation was found for SSRT_STOP_/SSRT_CS_ (*Pearson’s r_(49)_* = .41, *p_FDR_* = .003, CI_95_[.15, .61]) and SSRT_STOP_/SSRT_CSF_ (*r_(49)_* = .44, *p_FDR_* = .02, CI_95_[.19, .64]) while the SSRT_CS_/SSRT_CSF_ correlation was stronger (*r_(49)_* = .63, *p_FDR_* < .001, CI_95_[.43, .77]). This finding contrasts with the behavioral finding of Experiment 1, where there was no difference between SSRT_STOP_ and SSRT_CHANGE_. This is likely attributable to the differences in task design (see Discussion).

Similar to Experiment 1, response changing was faster than its primary initiation. GO-RT (M = 418 ms, SD = 57 ms) was longer than GO2_CS_-RT (M = 348 ms, SD = 90; *t_(50)_* = 6.70, *p_FDR_* < .001, CI_95_[50 ms, 92 ms], *d* = 0.85) and GO2_CSF_-RT (M = 332 ms, SD = 94 ms; *t_(50)_* = 7.90, *p_FDR_* < .001, CI_95_[64 ms, 108 ms], *d* = 0.97). Interestingly, GO2_CS_-RT was longer than GO2_CSF_-RT (*t_(50)_* = 3.31, *p_FDR_* = .002, CI_95_[6 ms, 25 ms], *d* = 0.17).

#### prEMG

Grand-average EMG waveforms and analysis results are depicted in **Figure 5** for Experiments 2 and 3. In addition to the participant excluded based on behavior (see above), 6 participants were excluded due to noisy EMG data (see method section for detailed criteria), resulting in a final sample of N = 45 analyzed datasets. Our manipulation-check procedure showed that EMG accurately captured the participants’ motor responses. The single-trial correlation between GO-RT and EMG peak latency was high for every channel (*Pearson’s r*, M = .93, SD = .07). Among 180 recorded channels (45 participants × 4 fingers), 8 were rejected, and an average 7% of trials were rejected in the retained channels. The proportion of prEMG identified across successfully aborted initial-responses was similar to previous investigations with stop-signal tasks (see Raud et al., 2022, for a meta-analysis). This proportion in SS trials (M = .42) was lower than SCS (M = .55, *t_(44)_* = 6.46, *p_FDR_* < .001, CI_95_[.09, .16], *d* = 0.68) and SCSF trials (M = .54, *t_(44)_* = 6.09, *p_FDR_* < .001, CI_95_[.08, .16], *d* = 0.63), while it did not differ between SCS and SCSF trials (*t_(44)_* = 0.77, *p* = .44, BF_01_ = 4.66).

**Figure 5.**
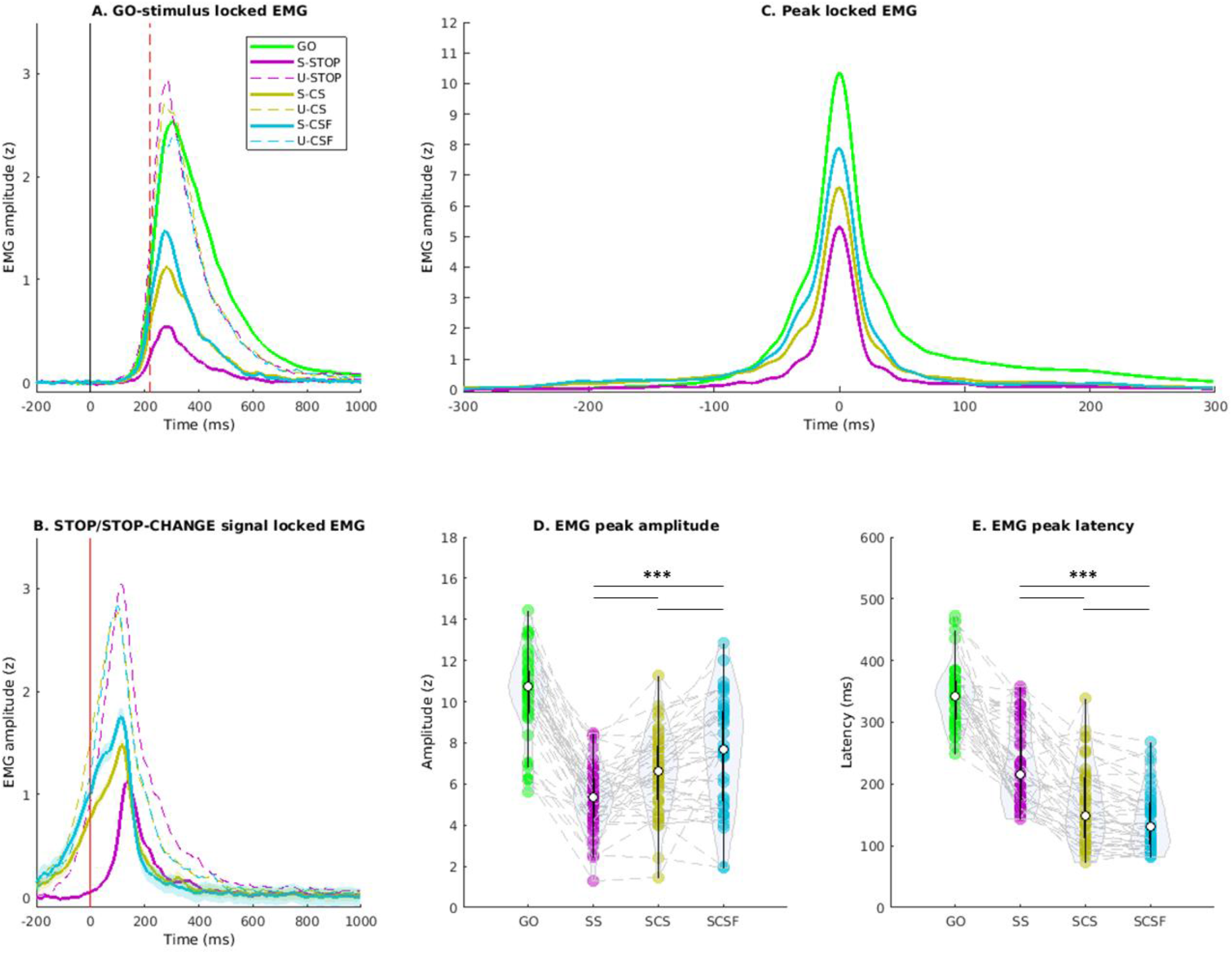
prEMG results of Experiment 2 & 3. Panel A, GO-stimulus locked grand-average EMG waveforms of the response triggered by the primary target (red line is the stop-signal occurrence time). Panel B, STOP/STOP-CHANGE signal locked grand-average EMG waveforms. Panel C, peak locked grand-average EMG waveforms. Panel D, individuals’ prEMG peak amplitude (and latency, panel E) for successfully aborted responses in SS, SCS and SCSF trials. \**p_FDR_* < .05, ** *p_FDR_* < .01, *** *p_FDR_* < .001.

Averaged waveforms (**Figure 5**) show that successfully aborted initial responses in SS, SCS and SCSF trials were associated with an EMG activation that was present to a lower extent than in response trials (GO, US, UCS, USF). In successfully aborted response, the amplitude of the prEMG (i.e., EMG activity that escaped inhibition) scaled from SS to SCS to SCSF. Indeed, prEMG peak amplitude was significantly lower in SS than SCS (*t_(44)_* = 5.02, *p_FDR_* < .001, CI_95_[0.77, 1.80 (z)], *d* = 0.68) and SCSF trials (*t_(44)_* = 5.53, *p_FDR_* < .001, CI_95_[1.48, 3.17 (z)], *d* = 0.95), while SCS prEMG amplitude was also lower than SCSF trials (*t_(44)_* = 3.81, *p_FDR_* < .001, CI_95_[0.49, 1.59 (z)], *d* = 0.44). Our results clearly show the presence of a prEMG in successful action-stopping, in line with previous studies (Raud et al., 2022). In addition, our analysis shows a scaling dynamic of the prEMG across our conditions, indicating that initial-response prEMG amplitude was higher in action-changing than -stopping. This suggests that additional motoric activity escaped inhibition in the aborted initial response when increasing the amount of required response revision.

The same analysis was performed for prEMG peak latency, showing that prEMG peak occurred earlier in CHANGE than STOP trials. Indeed, prEMG peak latency was later in SS (M = 230 ms, SD = 65 ms) than SCS (M = 165 ms, SD = 62 ms, *t_(44)_* = 6.63, *p_FDR_* < .001, CI_95_[45 ms, 85 ms], *d* = 0.91) and SCSF trials (M = 143 ms, SD = 50 ms, *t_(44)_* = 10.51, *p_FDR_* < .001, CI_95_[72 ms, 106 ms], *d* = 1.20). SCS and SCSF prEMG peak latencies also differ (*t_(44)_* = 4.44, *p_FDR_* < .001, CI_95_[10 ms, 26 ms], *d* = 0.39). These analyses showed a scaling dynamic across our conditions, the aborted initial-response prEMG peak being earlier when the amount of required response-revision was higher.

#### EEG

STOP/STOP-CHANGE signals induced a frontocentral β-burst dynamics modulation, especially visible as a β-burst volume increase (**Figure 6**). SSRT-window analysis revealed no significant modulation of the β-burst rate between STOP/STOP-CHANGE and match GO trials (*p_FDR_* > .05), but showed an increase of β-burst volume in SS (*t_(49)_* = 2.35, *p_FDR_* = .02, CI_95_[0.04, 0.51], *d* = 0.40), SCS (*t_(49)_* = 2.17, *p_FDR_*= .03, CI_95_[0.01, 0.31], *d* = 0.35), and SCSF trials (*t_(49)_* = 2.06, *p_FDR_* = .04, CI_95_[0.01, 0.44], *d* = 0.35), as compared to match GO trials. SSRT-window β-burst rate and volume did not show any significant difference across SSRT_STOP_, SSRT_CS,_ and SSRT_CSF_ (*p_FDR_* > .05, BF_01_ > 3). Overall, the β-burst volume showed an increase during SSRT which was identical across conditions. These results align with previous reports of post-STOP-signal frontocentral β-bursts dynamics (Enz et al., 2021) and support that β-bursts reflect the triggering of the ‘pause’ stage of inhibition (Tatz et al., 2023).

**Figure 6.**
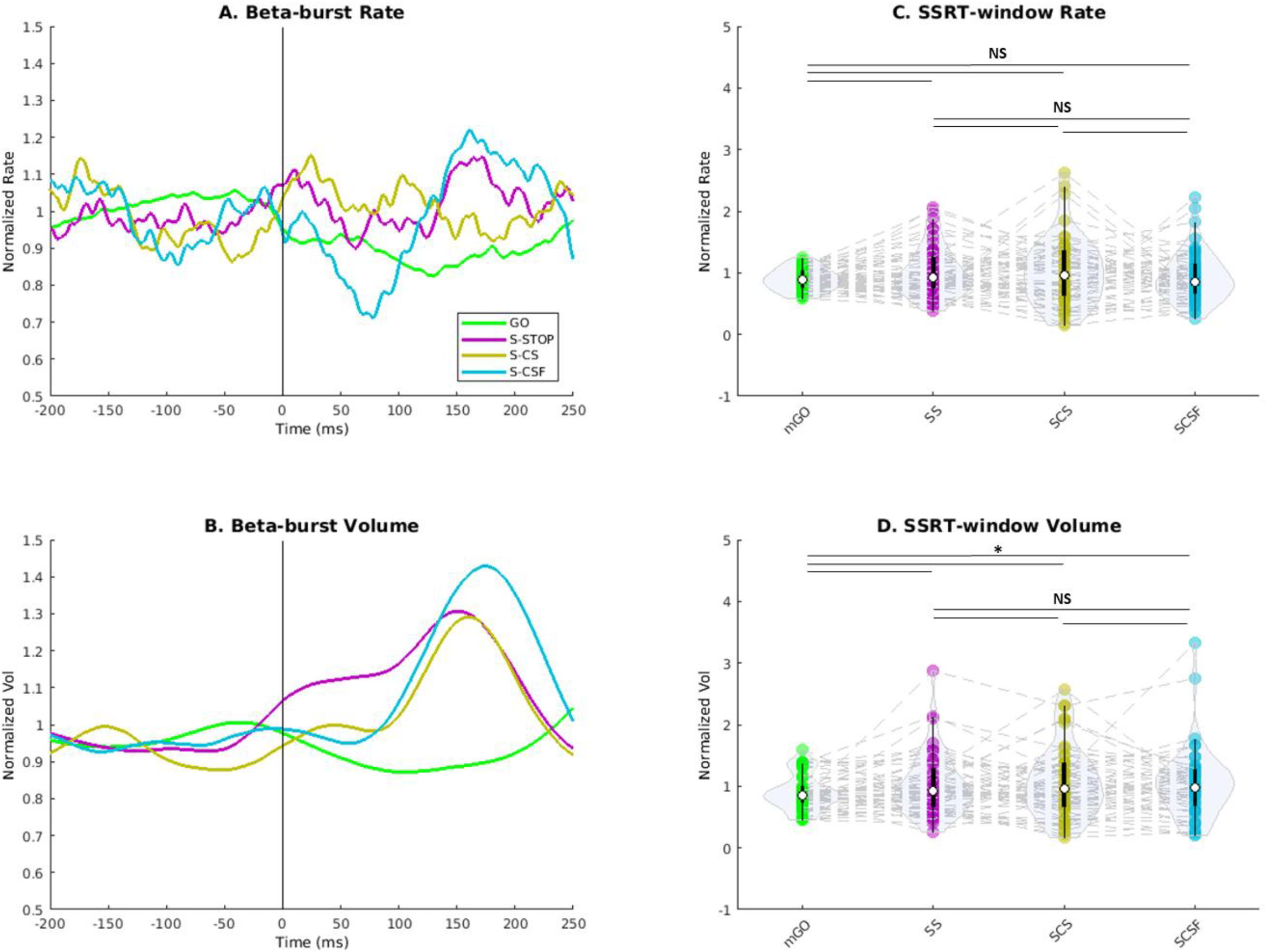
β-bursts results of Experiment 2 & 3. Panel A and B, grand-average β-burst rate and volume computed at FCz site and locked to the STOP/STOP-CHANGE signal. Panel C and D, individual’s averaged β-burst rate and volume within SSRT windows. \**p_FDR_* < .05, ** *p_FDR_* < .01, *** *p_FDR_* < .001.

ERPs showed a clear P3 wave evoked by the STOP/STOP-CHANGE signal with a frontocentral topography (**Figure 7**). In line with the visual inspection of the ERPs, the P3 onset analysis confirmed that the P3 wave was engaged in action-stopping and -changing as its onset was earlier in successfully compared to unsuccessfully canceled responses. Indeed, SS P3 onset (M = 247 ms, SD = 43 ms) was earlier than US (M = 274 ms, SD = 40 ms, one-tail paired *t*-test, *t_(40)_* = 3.70, *p_FDR_* < .001, CI_95_[14 ms, +∞], *d* = 0.62), this was also the case for SCS (M = 216 ms, SD = 54 ms) compared to USC (M = 250 ms, SD = 39 ms, *t_(41)_* = 3.71, *p_FDR_* < .001, CI_95_[16 ms, +∞], *d* = 0.67) and SCSF (M = 218 ms, SD = 37 ms) compared to UCSF (M = 239 ms, SD = 38 ms, *t_(44)_* = 5.25, *p_FDR_* < .001, CI_95_[15 ms, +∞], *d* = 0.52). These results confirmed the earlier P3 engagement in successful vs. unsuccessful action-stopping (Kok et al., 2004; Wessel and Aron, 2015) and expand the validity of this neurophysiological inhibition signature to action-changing. In line with our PTC model-based hypothesis, we observed that P3 amplitude scaled with the engaged action-revision. First, P3 amplitude was higher in successful changing conditions, compared to stopping. Second, P3 amplitude increased when adding a response dimension to revise (change-side-and-finger vs. change-side). Indeed, the P3 amplitude was lower in SS than SCS (one-tail paired *t*-test, *t_(40)_* = 2.30, *p_FDR_* = .016, CI_95_[0.20, +∞], *d* = 0.17) and SCSF trials (*t_(40)_* = 3.12, *p_FDR_* = .005, CI_95_[0.57, +∞], *d* = 0.26). Importantly, P3 was also larger in SCSF compared to SCS (*t_(42)_* = 2.21, *p_FDR_*= .016, CI_95_[0.12, +∞], *d* = 0.10), meaning that an additional response dimension to revise was associated with larger P3. We also computed an additive model to compare STOP-CHANGE P3 peak vs. [GO P3 + SS P3] peak. STOP-CHANGE P3 was still significantly higher for both SCS and SCSF conditions (*p_FDR_* < .05). Together, these results coherently indicate that frontocentral P3 wave scaled with the amount of engaged response revision, in a SS → SCS → SCSF order. Accordingly, θ power was increased post-STOP/STOP-CHANGE signal and scaled with the amount of response update. Indeed, SS θ power amplitude was lower than SCS (*t_(49)_* = 2.59, *p_FDR_* = .009, CI_95_[2.70, +∞], *d* = 0.23) and SCSF trials (*t_(49)_* = 2.79, *p_FDR_* = .009, CI_95_[2.87, +∞], *d* = 0.23). No significant difference between SCS and SCSF conditions (*t_(49)_* = 0.19, p_FDR_ = .57, BF_01_ = 7.50). Aligning with the P3 dynamic, these results showed that the θ power increase induced by a STOP-CHANGE signal was larger than a STOP signal. Though, the SCS vs. SCSF difference being unique to P3 might suggest that θ more generically reflects the larger cognitive control required in changing than stopping. In contrast, the finer P3 scaling according to the required response revision would reflect the actual ‘retuning’ of the motor response.

**Figure 7.**
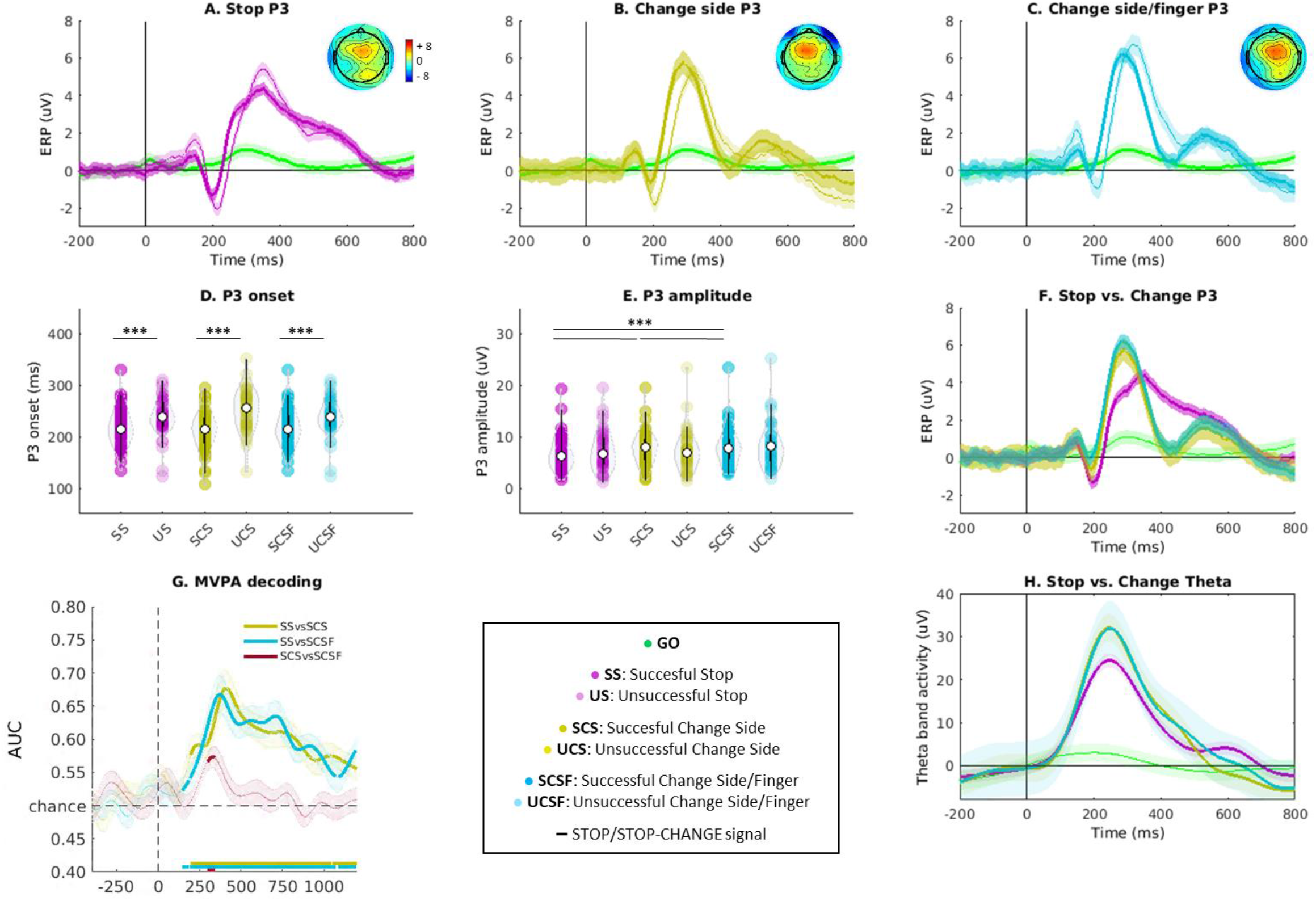
EEG results of Experiment 2 & 3. Panels A to C (and F), STOP/STOP-CHANGE-signal locked frontocentral grand-average ERP waveforms. Grand-average topographies were computed at the P3 peak latency identified at the group level. Panels D and E, individual’s P3 onset and amplitude. Panel G, whole-scalp EEG MVPA decoding performances. Horizontal colored lines indicate significant data points (see method). Panel H, frontocentral θ power time course (grand-average). \**p_FDR_* < .05, ** *p_FDR_* < .01, *** *p_FDR_* < .001.

Finally, the whole-scalp EEG MVPA decoding significantly decoded STOP and STOP-CHANGE trials starting 189 ms for SCS and 132 ms for SCSF. Maximum classification performances reached .68 (SS vs. SCS) and .67 (SS vs. SCSF). The MVPA procedure also succeeded in significantly distinguishing SCS and SCSF conditions starting at 311 ms (maximum performance of .58) – i.e., the exact time range of the frontocentral P3. These results suggest that STOP and CHANGE conditions started to engage unique EEG activity in an early post-signal window compatible with the late-edge of the proposed ‘pause’ inhibitory stage. STOP-CHANGE conditions began to differ, depending on the number of response dimensions (side vs. side-and-finger), in a later window compatible with the assumed ‘cancel’/’retune’ timeline.

## Discussion

Past work on action-stopping has identified several candidate signatures of inhibitory control. However, they have been hard to reconcile under a joint framework, owing largely to their timing differences and debated functional significance. Evaluating these signatures altogether for the first time, we successfully disentangle their functional properties across multiple action-revision conditions. Accounting for common and unique neurophysiological dynamics, our results support a ‘pause-then-retune’ conceptualization of human action-revision in line with two-stage models of action-stopping (Schmidt and Berke, 2017; Diesburg and Wessel, 2021). Early signatures (non-selective CSE suppression and β-burst volume increase) were common to stopping and changing, in line with a shared ‘pause’-stage inhibition processing. In contrast, later frontocentral θ power was larger for changing than stopping, and P3 scaled with the amount of response revision, in line with the proposal that these signatures reflect a separate ‘retuning’ stage, which is sensitive to the exact demands of the task.

Our results have several notable features. First, the early non-selective CSE suppression was present in both action-stopping and -changing. This non-selective CSE suppression has been causally associated with the subthalamic nucleus (STN) in action-stopping based on TMS/DBS combination (Wessel et al., 2022). Therefore, our result is congruent with previous findings from fMRI (Kenner et al., 2010) and STN-DBS in Parkinson’s disease (van den Wildenberg et al., 2017, 2021) supporting the engagement of the STN-mediated hyperdirect pathway in both action-stopping and -changing. Similar early CSE suppression corroborated our prediction that broad inhibition is commonly triggered by a ‘pause’ process in the two situations, and rejects the alternative view (Logan and Burkell, 1986; De Jong et al., 1995; Krämer et al., 2011) of inhibition being mainly implemented by “*selectively preventing central outflow of motor commands*” in action-changing context (De Jong et al., 1995). Besides, increases in SSRT-window β-burst volume did not differ across action-revision conditions, with moderate evidence supporting the null hypothesis. This might suggest that directional subcortical-cortical β-bursting associated with the hyperdirect pathway (Torrecillos et al., 2018; Diesburg et al., 2021) is part of a ‘pause’ process that triggers early broad inhibition in a fully non-selective fashion. This speculative proposal would require a more direct investigation of the subcortical dynamic within action-revision, including intracranial recording from the STN.

Second, prEMG showed that some motor activation from the aborted initial response survived the ‘pause’-stage inhibition. This general finding speaks towards the fact that action-stopping may not be carried out by a single-process. Our specific hypothesis in the current study was that we would find common prEMG dynamics across action-revision conditions, reflecting a shared ‘pause’ process. In contrast, prEMG trials count, amplitude, and timing scaled across our conditions, indicating that more residual motoric activity escaped inhibition in change-side-and-finger than in change-side and then stop successful trials. As the EMG reflects a net output of the motor system, its properties might actually reflect both the engagement of the ‘pause’ and ‘cancel’ (or ‘retune’, see below) processes (Fisher et al., 2024), depending on the task demand. Accordingly, the broad motor inhibition triggered early in both stopping and changing by the ‘pause’ process (reflected in early-CSE and β-bursts) might then be sustained (in stopping) or released (in changing) depending on the required revision. This release might thus have led to earlier/larger prEMG in action-changing. Presumably, the broad-inhibition release facilitated the redirection of the initial motor activation to speed its ‘retuning’ and, hence, the revised response (as reflected by prEMG/GO2-RTs correlations and faster GO2-RTs than initial GO-RT). Recent proposals that the ‘pause’ vs. ‘cancel’ relative timing varies based on individuals’ strategies and task context (Hannah et al., 2022; Pani et al., 2022) might suggest that this quick redirection of the motor activity differs across our experiments. Indeed, a surprising result was that SSRT was much faster for changing than stopping in Experiments 2 and 3, which was not the case in Experiment 1. Although SSRT has been widely criticized as a latent construct (Boucher et al., 2007; Skippen et al., 2019; Huster et al., 2020; Bissett et al., 2021), this difference could reflect a task-design effect on the speed of redirecting the motor command. Indeed, GO and STOP-CHANGE stimuli instructions were associated with a specific symbol (left vs. right arrow) in Experiment 1, while they relied on tracking the green target in Experiments 2 and 3. Motor redirection from the aborted initial response to the new one might be speeded in this tracking context. Further investigation using multiple effectors could address this hypothesis, as various effector combinations have been associated with modulations of stopping vs. changing SSRT differences (Boecker et al., 2013). Dual-coil TMS assessment of CSE might also provide further insight into this proposed quick transfer of the motor command (Ratcliffe et al., 2022; Wadsley et al., 2023b).

Third, a frontocentral P3 ERP was observed in both action-stopping and -changing, alongside an earlier P3 onset latency between successfully vs. unsuccessfully canceled responses. Beyond the replication of P3 decisive role in successful action-stopping (Kok et al., 2004; Wessel and Aron, 2015; Dykstra et al., 2020; Tatz et al., 2021), Experiments 2 and 3 showed that P3 peak amplitude consistently increased according to the required response revision in an SS → SCS → SCSF order. The maximal decoding performances of the whole-scalp EEG MVPA fell in the same time range, including for the SCS vs. SCSF contrast, and P3_CHANGE_ was higher than P3_STOP+GO_. These support further that the P3 was specifically increased in action-changing compared to -stopping, and in changing two response dimensions compared to a single one. Following the view of inhibition being implemented similarly between action-stopping and -changing [Change = Stop + Reengage] (Boecker et al., 2013), one might expect that the ‘cancel’ stage, and P3 as its purported signature, completes the initial-response suppression independently of the new response engagement, though overlapping in time (Verbruggen et al., 2008). In contrast, the P3 modulation we reported may reflect the need for additional cognitive control in action-changing. Based on a ‘context-updating-theory’ of P3-type waveforms (Donchin and Coles, 1988; Polich, 2007), the larger update required in the motor/task representation generated by the primary stimulus could be associated with an increased P3 in action-changing. Higher frontocentral θ power evoked by the STOP-CHANGE vs. STOP signal may also pair with higher cognitive demand (Huster et al., 2013; Messel et al., 2021). Therefore, we propose that the revision could be performed by a selective ‘retuning’ of the motor command that results in canceling or redirecting the motoric activity that survived the ‘pause’ inhibition, depending on the task requirements. In favor of this view, previous studies have reported an increased P3 amplitude when an infrequent signal instructed stopping an action compared to continuing it (Waller et al., 2019; Tatz et al., 2021; González-Villar et al., 2022). Our new finding of P3 amplitudes that reflect the amount of response revision is in line with the view of a ‘pause-then-retune’ dynamic triggered by any infrequent event (Wessel and Aron, 2017), the ‘retune’-stage neurophysiological activity scaling with the required motor-command revision: in the current case, either continuing, stopping, or changing the response. To specify how the motor response retuning is implemented beyond our discrete approach (change-side vs. change-side-and-finger), future work might modulate the amount of engaged response change in a continuous fashion, e.g., based on movement trajectory tracking (Zanone and Athènes, 2013; Erb et al., 2016; Hervault et al., 2019; Temprado et al., 2020; London et al., 2021). Moreover, such specific ‘retuning’ of the motor command might specifically engage the pre-SMA, whose role has been previously suggested in reweighting action programs (Jha et al., 2015; Diesburg and Tatz, 2021; Hervault et al., 2022; Wolpe et al., 2022). Although it is still speculative at this stage, this proposal would align with the finding that pre-SMA lesions impaired action-changing to a larger extent than -stopping (Roberts and Husain, 2015).

In conclusion, we aimed to assess the neurophysiological signatures of human action-revision in a previously unseen, comprehensive fashion. Combining EEG, EMG, TMS, and a novel experimental paradigm led to unique insights into the differential dynamics of action-stopping and action-changing. Common and unique neurophysiological processes between action-stopping and -changing led us to propose that inhibitory control is implemented in a generic ‘pause-then-retune’ dynamic that specifies its mechanism depending on the required response revision.

## Acknowledgments

This work was supported by National Science Foundation Career Grant 1752355 to J.R.W.; and National Institutes of Health Grant R01 NS117753 to J.R.W. The authors would like to thank all the volunteers for their participation, Nathan Chalkley, Nathan Chen and Saara Engineer for help in data acquisition.

## Author contributions

M.H. and J.R.W. designed research; M.H. performed research; M.H. analyzed data; M.H. and J.R.W. edited the paper; M.H. and J.R.W. wrote the paper; M.H. wrote the first draft of the paper.

## Bibliography

Antons S, Boecker M, Gauggel S, Gordi VM, Patel HJ, Binkofski F, Drueke B (2019) Strategies of selective changing: Preparatory neural processes seem to be responsible for differences in complex inhibition. PLOS ONE 14:e0214652.

Badry R, Mima T, Aso T, Nakatsuka M, Abe M, Fathi D, Foly N, Nagiub H, Nagamine T, Fukuyama H (2009) Suppression of human cortico-motoneuronal excitability during the Stop-signal task. Clinical Neurophysiology 120:1717–1723.

Bell AJ, Sejnowski TJ (1995) An information-maximization approach to blind separation and blind deconvolution. Neural Comput 7:1129–1159.

Benjamini Y, Krieger AM, Yekutieli D (2006) Adaptive linear step-up procedures that control the false discovery rate. Biometrika 93:491–507.

Bissett PG, Jones HM, Poldrack RA, Logan GD (2021) Severe violations of independence in response inhibition tasks. Science Advances.

Bissett PG, Logan GD (2014) Selective stopping? Maybe not. J Exp Psychol Gen 143:455–472.

Boecker M, Buecheler MM, Schroeter ML, Gauggel S (2007) Prefrontal brain activation during stop-signal response inhibition: An event-related functional near-infrared spectroscopy study. Behavioural Brain Research 176:259–266.

Boecker M, Drueke B, Vorhold V, Knops A, Philippen B, Gauggel S (2011) When response inhibition is followed by response reengagement: An event-related fMRI study. Human Brain Mapping 32:94–106.

Boecker M, Gauggel S, Drueke B (2013) Stop or stop-change — Does it make any difference for the inhibition process? International Journal of Psychophysiology 87:234–243.

Boucher L, Palmeri TJ, Logan GD, Schall JD (2007) Inhibitory control in mind and brain: an interactive race model of countermanding saccades. Psychol Rev 114:376–397.

Brainard DH (1997) The Psychophysics Toolbox. Spatial Vision 10:433–436.

Cagnan H, Mallet N, Moll CKE, Gulberti A, Holt AB, Westphal M, Gerloff C, Engel AK, Hamel W, Magill PJ, Brown P, Sharott A (2019) Temporal evolution of beta bursts in the parkinsonian cortical and basal ganglia network. Proceedings of the National Academy of Sciences 116:16095–16104.

Choo Y, Matzke D, Bowren MD Jr, Tranel D, Wessel JR (2022) Right inferior frontal gyrus damage is associated with impaired initiation of inhibitory control, but not its implementation Badre D, Frank MJ, Fellows LK, eds. eLife 11:e79667.

De Jong R, Coles MGH, Logan GD (1995) Strategies and mechanisms in nonselective and selective inhibitory motor control. Journal of Experimental Psychology: Human Perception and Performance 21:498–511.

Delorme A, Makeig S (2004) EEGLAB: an open source toolbox for analysis of single-trial EEG dynamics including independent component analysis. J Neurosci Methods 134:9–21.

Diesburg DA, Greenlee JD, Wessel JR (2021) Cortico-subcortical β burst dynamics underlying movement cancellation in humans Swann NC, Ivry RB, Muralidharan V, Schmidt R, eds. eLife 10:e70270.

Diesburg DA, Tatz JR (2021) Unexpected Events Activate a Frontal-Basal-Ganglia Inhibitory Network: What Is the Role of the Pre-Supplementary Motor Area? J Neurosci 41:5135– 5137.

Diesburg DA, Wessel JR (2021) The Pause-then-Cancel model of human action-stopping: Theoretical considerations and empirical evidence. Neuroscience & Biobehavioral Reviews 129:17–34.

Donchin E, Coles MGH (1988) Is the P300 component a manifestation of context updating? Behavioral and Brain Sciences 11:357–374.

Drueke B, Boecker M, Schlaegel S, Moeller O, Hiemke C, Gründer G, Gauggel S (2010) Serotonergic modulation of response inhibition and re-engagement? Results of a study in healthy human volunteers. Human Psychopharmacology: Clinical and Experimental 25:472–480.

Duque J, Greenhouse I, Labruna L, Ivry RB (2017) Physiological Markers of Motor Inhibition during Human Behavior. Trends in Neurosciences 40:219–236.

Dykstra T, Waller DA, Hazeltine E, Wessel JR (2020) Leveling the Field for a Fairer Race between Going and Stopping: Neural Evidence for the Race Model of Motor Inhibition from a New Version of the Stop Signal Task. J Cogn Neurosci 32:590–602.

Elchlepp H, Verbruggen F (2017) How to withhold or replace a prepotent response: An analysis of the underlying control processes and their temporal dynamics. Biological Psychology 123:250–268.

Enz N, Ruddy KL, Rueda-Delgado LM, Whelan R (2021) Volume of β-Bursts, But Not Their Rate, Predicts Successful Response Inhibition. J Neurosci 41:5069–5079.

Erb CD, Moher J, Sobel DM, Song J-H (2016) Reach tracking reveals dissociable processes underlying cognitive control. Cognition 152:114–126.

Fahrenfort JJ, van Driel J, van Gaal S, Olivers CNL (2018) From ERPs to MVPA Using the Amsterdam Decoding and Modeling Toolbox (ADAM). Frontiers in Neuroscience 12.

Faul F, Erdfelder E, Buchner A, Lang A-G (2009) Statistical power analyses using G*Power 3.1: Tests for correlation and regression analyses. Behavior Research Methods 41:1149– 1160.

Fisher M, Trinh H, O’Neill J, Greenhouse I (2024) Early Rise and Persistent Inhibition of Electromyography during Failed Stopping. Journal of Cognitive Neuroscience:1–15.

Giarrocco F, Bardella G, Giamundo M, Fabbrini F, Brunamonti E, Pani P, Ferraina S (2021) Neuronal dynamics of signal selective motor plan cancellation in the macaque dorsal premotor cortex. Cortex 135:326–340.

González-Villar A, Galdo-Álvarez S, Carrillo-de-la-Peña MT (2022) Neural correlates of unpredictable Stop and non-Stop cues in overt and imagined execution. Psychophysiology 59:e14019.

Hannah R, Jana S, Muralidharan V (2022) Does action-stopping involve separate pause and cancel processes? A view from premotor cortex. Cortex 152:157–159.

Hannah R, Muralidharan V, Sundby KK, Aron AR (2020) Temporally-precise disruption of prefrontal cortex informed by the timing of beta bursts impairs human action-stopping. NeuroImage 222:117222.

Hervault M, Huys R, Farrer C, Buisson JC, Zanone PG (2019) Cancelling discrete and stopping ongoing rhythmic movements: Do they involve the same process of motor inhibition? Hum Mov Sci 64:296–306.

Hervault M, Zanone P-G, Buisson J-C, Huys R (2022) Multiple Brain Sources Are Differentially Engaged in the Inhibition of Distinct Action Types. Journal of Cognitive Neuroscience 34:258–272.

Huster RJ, Enriquez-Geppert S, Lavallee CF, Falkenstein M, Herrmann CS (2013) Electroencephalography of response inhibition tasks: functional networks and cognitive contributions. Int J Psychophysiol 87:217–233.

Huster RJ, Messel MS, Thunberg C, Raud L (2020) The P300 as marker of inhibitory control – Fact or fiction? Cortex.

Hynd M, Soh C, Rangel BO, Wessel JR (2021) Paired-pulse TMS and scalp EEG reveal systematic relationship between inhibitory GABAa signaling in M1 and fronto-central cortical activity during action stopping. Journal of Neurophysiology 125:648–660.

Jana S, Hannah R, Muralidharan V, Aron AR (2020) Temporal cascade of frontal, motor and muscle processes underlying human action-stopping van den Wildenberg W, Ivry RB, van den Wildenberg W, Huster R, Bissett PG, eds. eLife 9:e50371.

Jha A, Nachev P, Barnes G, Husain M, Brown P, Litvak V (2015) The Frontal Control of Stopping. Cereb Cortex 25:4392–4406.

Kenner NM, Mumford JA, Hommer RE, Skup M, Leibenluft E, Poldrack RA (2010) Inhibitory motor control in response stopping and response switching. J Neurosci 30:8512–8518.

Kok A, Ramautar JR, De Ruiter MB, Band GPH, Ridderinkhof KR (2004) ERP components associated with successful and unsuccessful stopping in a stop-signal task. Psychophysiology 41:9–20.

Krämer UM, Knight RT, Münte TF (2011) Electrophysiological evidence for different inhibitory mechanisms when stopping or changing a planned response. J Cogn Neurosci 23:2481– 2493.

Krekelberg B (2023) bayesFactor. Available at: https://github.com/klabhub/bayesFactor.

Logan GD, Burkell J (1986) Dependence and independence in responding to double stimulation: A comparison of stop, change, and dual-task paradigms. Journal of Experimental Psychology: Human Perception and Performance 12:549–563.

Logan GD, Cowan WB (1984) On the ability to inhibit thought and action: A theory of an act of control. Psychological Review 91:295–327.

London D, Fazl A, Katlowitz K, Soula M, Pourfar MH, Mogilner AY, Kiani R (2021) Distinct population code for movement kinematics and changes of ongoing movements in human subthalamic nucleus. eLife 10:e64893.

Majid DSA, Cai W, George JS, Verbruggen F, Aron AR (2012) Transcranial magnetic stimulation reveals dissociable mechanisms for global versus selective corticomotor suppression underlying the stopping of action. Cereb Cortex 22:363–371.

Maris E, Oostenveld R (2007) Nonparametric statistical testing of EEG- and MEG-data. J Neurosci Methods 164:177–190.

McClure EB, Treland JE, Snow J, Schmajuk M, Dickstein DP, Towbin KE, Charney DS, Pine DS, Leibenluft E (2005) Deficits in Social Cognition and Response Flexibility in Pediatric Bipolar Disorder. AJP 162:1644–1651.

Messel MS, Raud L, Hoff PK, Stubberud J, Huster RJ (2021) Frontal-midline theta reflects different mechanisms associated with proactive and reactive control of inhibition. NeuroImage 241:118400.

Pani P, Giarrocco F, Bardella G, Brunamonti E, Ferraina S (2022) Reply to: Hannah et al. (2021) Commentary: ‘Does action-stopping involve separate pause and cancel processes? A view from premotor cortex’: Action-stopping models must consider the role of the dorsal premotor cortex. Cortex 152:160–163.

Pion-Tonachini L, Kreutz-Delgado K, Makeig S (2019) ICLabel: An automated electroencephalographic independent component classifier, dataset, and website. Neuroimage 198:181–197.

Polich J (2007) Updating P300: An Integrative Theory of P3a and P3b. Clin Neurophysiol 118:2128–2148.

Rangel-Gomez M, Knight RT, Krämer UM (2015) How to stop or change a motor response: Laplacian and independent component analysis approach. International Journal of Psychophysiology 97:233–244.

Ratcliffe CE, McAllister CJ, MacDonald HJ (2022) Dual-Coil Transcranial Magnetic Stimulation Reveals Temporal Dynamics of Bilateral Corticomotor Excitability During Response Inhibition. :2022.09.14.507942 Available at: https://www.biorxiv.org/content/10.1101/2022.09.14.507942v1.

Raud L, Huster RJ, Ivry RB, Labruna L, Messel MS, Greenhouse I (2020) A Single Mechanism for Global and Selective Response Inhibition under the Influence of Motor Preparation. J Neurosci 40:7921–7935.

Raud L, Thunberg C, Huster RJ (2022) Partial response electromyography as a marker of action stopping Badre D, Frank MJ, Leunissen I, eds. eLife 11:e70332.

Roberts RE, Husain M (2015) A dissociation between stopping and switching actions following a lesion of the pre-supplementary motor area. Cortex 63:184–195.

Rossi S, Hallett M, Rossini PM, Pascual-Leone A (2011) Screening questionnaire before TMS: An update. Clinical Neurophysiology 122:1686.

Rossini PM et al. (2015) Non-invasive electrical and magnetic stimulation of the brain, spinal cord, roots and peripheral nerves: Basic principles and procedures for routine clinical and research application. An updated report from an I.F.C.N. Committee. Clinical Neurophysiology 126:1071–1107.

Schmidt R, Berke JD (2017) A Pause-then-Cancel model of stopping: evidence from basal ganglia neurophysiology. Philos Trans R Soc Lond B Biol Sci 372.

Skippen P, Fulham WR, Michie PT, Matzke D, Heathcote A, Karayanidis F (2020) Reconsidering electrophysiological markers of response inhibition in light of trigger failures in the stop-signal task. Psychophysiology 57:e13619.

Skippen P, Matzke D, Heathcote A, Fulham WR, Michie P, Karayanidis F (2019) Reliability of triggering inhibitory process is a better predictor of impulsivity than SSRT. Acta Psychologica 192:104–117.

Tatz JR, Mather A, Wessel JR (2023) β-Bursts over Frontal Cortex Track the Surprise of Unexpected Events in Auditory, Visual, and Tactile Modalities. Journal of Cognitive Neuroscience 35:485–508.

Tatz JR, Soh C, Wessel JR (2021) Common and unique inhibitory control signatures of action-stopping and attentional capture suggest that actions are stopped in two stages. J Neurosci.

Temprado J-J, Torre MM, Langeard A, Julien-Vintrou M, Devillers-Réolon L, Sleimen-Malkoun R, Berton E (2020) Intentional Switching Between Bimanual Coordination Patterns in Older Adults: Is It Mediated by Inhibition Processes? Frontiers in Aging Neuroscience 12.

Torrecillos F, Tinkhauser G, Fischer P, Green AL, Aziz TZ, Foltynie T, Limousin P, Zrinzo L, Ashkan K, Brown P, Tan H (2018) Modulation of Beta Bursts in the Subthalamic Nucleus Predicts Motor Performance. J Neurosci 38:8905–8917.

van den Wildenberg WPM, Ridderinkhof KR, van Wouwe NC, Neimat JS, Bashore TR, Wylie SA (2017) Overriding actions in Parkinson’s disease: Impaired stopping and changing of motor responses. Behavioral Neuroscience 131:372–384.

van den Wildenberg WPM, van Wouwe NC, Ridderinkhof KR, Neimat JS, Elias WJ, Bashore TR, Wylie SA (2021) Deep-brain stimulation of the subthalamic nucleus improves overriding motor actions in Parkinson’s disease. Behavioural Brain Research 402:113124.

Verbruggen F et al. (2019) A consensus guide to capturing the ability to inhibit actions and impulsive behaviors in the stop-signal task Frank MJ, Badre D, Egner T, Swick D, eds. eLife 8:e46323.

Verbruggen F, Logan GD (2009) Models of response inhibition in the stop-signal and stop-change paradigms. Neurosci Biobehav Rev 33:647–661.

Verbruggen F, Schneider DW, Logan GD (2008) How to stop and change a response: the role of goal activation in multitasking. J Exp Psychol Hum Percept Perform 34:1212–1228.

Wadsley CG, Cirillo J, Nieuwenhuys A, Byblow WD (2023a) A global pause generates nonselective response inhibition during selective stopping. Cerebral Cortex 33:9729– 9740.

Wadsley CG, Cirillo J, Nieuwenhuys A, Byblow WD (2023b) Proactive Interhemispheric Disinhibition Supports Response Preparation during Selective Stopping. J Neurosci 43:1008–1017.

Waller DA, Hazeltine E, Wessel JR (2019) Common neural processes during action-stopping and infrequent stimulus detection: The frontocentral P3 as an index of generic motor inhibition. International Journal of Psychophysiology.

Wessel JR (2020) β-Bursts Reveal the Trial-to-Trial Dynamics of Movement Initiation and Cancellation. J Neurosci 40:411–423.

Wessel JR, Aron AR (2015) It’s not too late: the onset of the frontocentral P3 indexes successful response inhibition in the stop-signal paradigm. Psychophysiology 52:472–480.

Wessel JR, Aron AR (2017) On the Globality of Motor Suppression: Unexpected Events and Their Influence on Behavior and Cognition. Neuron 93:259–280.

Wessel JR, Diesburg DA, Chalkley NH, Greenlee JDW (2022) A causal role for the human subthalamic nucleus in non-selective cortico-motor inhibition. Current Biology 32:3785–3791.e3.

Wessel JR, Reynoso HS, Aron AR (2013) Saccade suppression exerts global effects on the motor system. Journal of Neurophysiology 110:883–890.

Wickens TD (2002) Elementary signal detection theory. New York, NY, US: Oxford University Press.

Wolpe N, Hezemans FH, Rae CL, Zhang J, Rowe JB (2022) The pre-supplementary motor area achieves inhibitory control by modulating response thresholds. Cortex 152:98–108.

Zanone P-G, Athènes S (2013) Switching among graphic patterns is governed by oscillatory coordination dynamics: implications for understanding handwriting. Front Psychol 4:662.

